# Disrupted RNA editing in beta cells mimics early stage type 1 diabetes

**DOI:** 10.1101/2022.12.08.519618

**Authors:** Udi Ehud Knebel, Shani Peleg, Chunhua Dai, Roni Cohen-Fultheim, Benjamin Glaser, Erez Y. Levanon, Alvin C. Powers, Agnes Klochendler, Yuval Dor

**Author notes:** Correspondence: Yuval Dor, or Agnes Klochendler. Equal contributors (Udi Ehud Knebel, Shani Peleg).

## Abstract

A major hypothesis for the etiology of type 1 diabetes (T1D) postulates initiation by viral infection, leading to double-stranded RNA (dsRNA)-mediated interferon response; however, a causal virus has not been identified. Here we use a mouse model, corroborated with human data, to demonstrate that endogenous dsRNA in beta-cells can lead to a diabetogenic immune response, thus identifying a virus-independent mechanism for T1D initiation. We found that disruption of the RNA editing enzyme ADAR in beta-cells triggers a massive interferon response, islet inflammation and beta-cell failure, with features bearing striking similarity to early-stage human T1D. Glycolysis via calcium enhances the interferon response, suggesting an actionable vicious cycle of inflammation and increased beta-cell workload.

**One sentence summary:** Adar inactivation in beta-cells triggers a glucose-dependent interferon response causing insulitis and diabetes

---

Adenosine to Inosine (A-to-I) RNA editing is a post-transcriptional modification catalyzed by Adenosine Deaminases Acting on RNA (ADAR), which deaminate adenosine bases into inosines, read by the translation machinery as guanosine (*1, 2*). Recent studies indicate that a key role of ADAR1 is to edit double-stranded RNA (dsRNA) structures, primarily generated by retroelements (typically *Alu* and B1/B2 repeats in human and mouse cells, respectively) inserted in an inverted orientation in non-coding regions of expressed genes (*2*). Such dsRNA structures are powerful and potentially dangerous activators of dsRNA sensors, such as MDA5/IFIH1 that triggers an antiviral type I interferon (IFN-I) response (*2*). Thus ADAR1-mediated A-to-I RNA editing destabilizes A-U base pairing in RNA, preventing pathogenic activation of an interferon response triggered by endogenous dsRNA. Indeed, ADAR1 mutations in humans and mice cause severe auto-inflammatory disease (*3, 4*).

While RNA editing has been studied in several organs in mouse models and in cancer (*5–8*), little is known about its role in insulin-producing beta-cells. This is particularly relevant since early stages of type I diabetes (T1D), an autoimmune disease causing the destruction of beta-cells, are associated with an IFN-I transcriptional signature in pancreatic islets (*9–12*). In addition, GWAS studies have shown that *IFIH1* variants strongly modulate T1D risk, suggesting the importance of sensing dsRNA – from either a viral or endogenous source – for the development of T1D (*13, 14*). A viral infection in islets could lead to an interferon response but such a virus has not been identified (*15, 16*). In support of endogenous dsRNA being a T1D trigger, whole-body loss of *Ifih1* was shown to prevent spontaneous autoimmune diabetes in NOD mice, independently of viral infection (*17*). Furthermore, a recent study reported that human genetic variants reducing RNA editing of endogenous dsRNAs confer increased risk of autoinflammatory and autoimmune diseases, including T1D (*18*). We hypothesized that disruption of Adar in beta-cells *in vivo* may model early events in the pathogenesis of T1D and allow dissection of mechanisms by which non-viral, endogenous dsRNA can trigger islet inflammation and contribute to beta-cell dysfunction or loss, prior to autoimmunity.

To test these ideas, we disrupted *Adar* in beta-cells of mice and knocked down *ADAR* in human islets. We show that *ADAR* inactivation in beta-cells elicits a strong interferon response, leading *in vivo* to islet infiltration and impairment of beta-cell function and survival. We further delineate cell-autonomous and paracrine determinants of beta-cell failure following *Adar* inactivation, and describe phenotypes that strikingly resemble early stage T1D. Finally, we reveal a modulation of the beta-cell interferon response by glucose metabolism and calcium signaling, suggesting an actionable positive feedback loop whereby increased beta-cell workload and islet inflammation drive beta-cell failure.

## Results

Most RNA editing events in mammals are catalyzed by ADAR1 (Adar), and map to pairs of inverted repetitive elements in introns or UTRs of transcripts that can form dsRNA structures (*19, 20*). The extent of editing correlates with the number of repetitive elements within families with low sequence divergence (*21–23*). Consistently, the *Alu* editing index in human cells is higher than the editing index in the more divergent mouse B1/B2 repeats (*24*).

We applied our *Alu* Editing Index (AEI), which provides information about the general level of editing in repeats, to published RNA-Seq datasets from human and mouse alpha- and beta-cells to quantify highly edited regions, and particularly extensively edited regions in 3’-UTR inverted repeats (*25*). As expected, A-to-G RNA editing index was enriched compared to the index of other mismatches indicating highly specific detection of ADAR-mediated editing. Human alpha- and beta-cells had a higher A-to-G RNA editing index levels compared with mouse cells, while in both species there were no significant differences between editing in alpha- and beta-cells (**Fig. 1A-B**).

**Fig.1:**
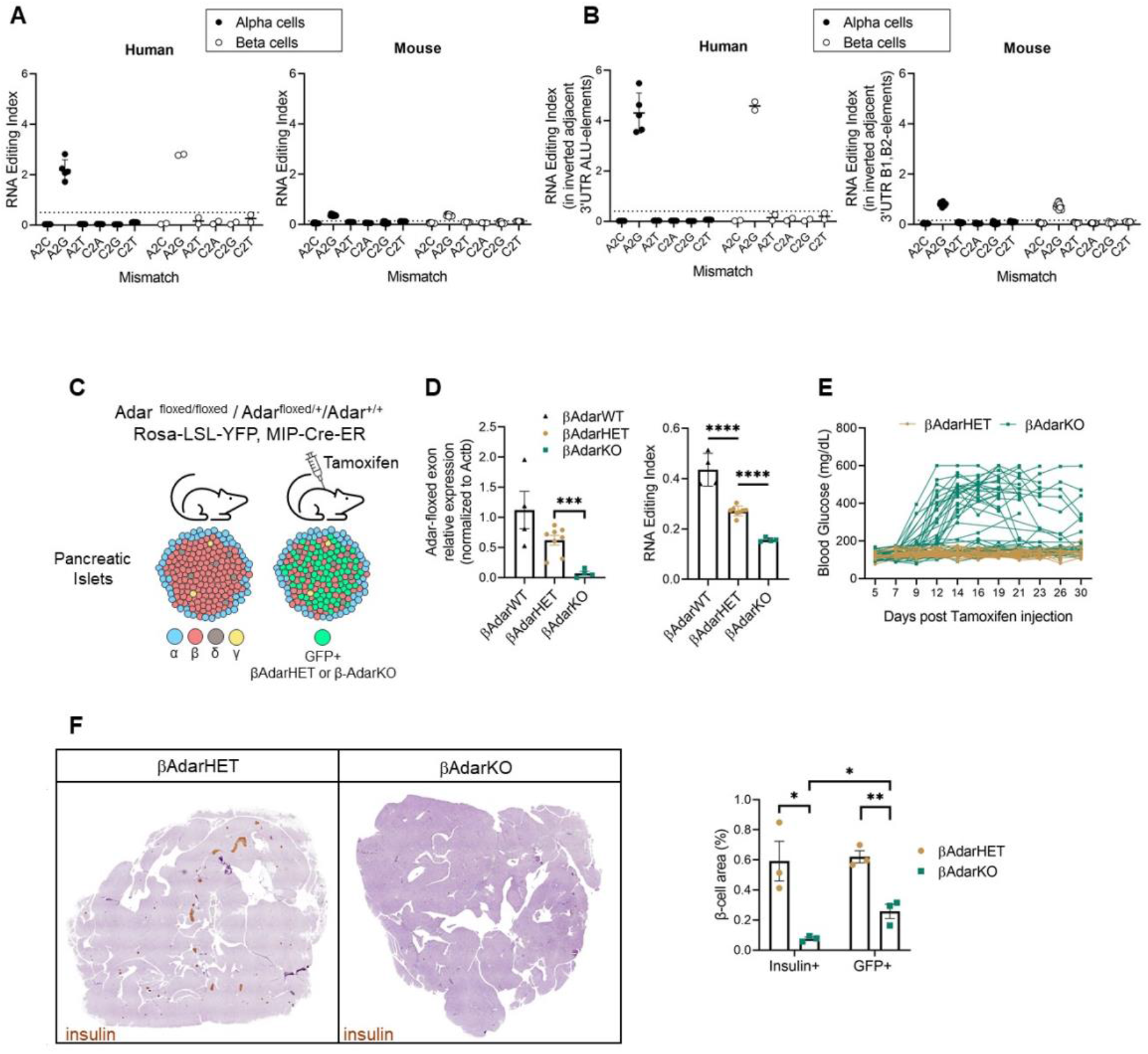
Loss of RNA editing in mouse beta-cells disrupts glucose homeostasis. (**A**) Global A-to-I RNA editing index (*25*) across short interspersed nuclear elements (SINE) in human (*Alu*) and mouse (B1 and B2) RNA-seq data (*53–59*) demonstrates a higher A-to-I editing signal in human samples. Noise levels (non-A-to-G mismatches) are similar between human and mouse. (**B**) A-to-I RNA editing in inverted elements within 3’UTR is twice as high as the global editing index. Noise levels remain similar or lower than seen in the global editing index. Minimal element length was set to reflect average element length in human and mouse (200bp and 120bp, respectively). (**C**) Experimental approach to study the effects of Adar inactivation in mouse beta-cells. (**D**) Reduced expression of Adar floxed exons and reduced editing at oppositely oriented B1/B2 repetitive elements in RNA from βAdarKO versus βAdarHET and βAdarWT beta-cells. Data are means ± SE. Student’s unpaired two-tailed t-test; *** P<0.001, **** P<0.0001 (**E**) Blood glucose was measured for 30 days after Tamoxifen injection in βAdarHET (n=36). and βAdarKO (n=52) mice (N=27 males, 25 females). (**F**) Representative micrographs of pancreatic sections from βAdarHET and diabetic βAdarKO mice 1 month after tamoxifen injection, immunostained for Insulin. Insulin and GFP positive area per pancreas were measured 1 month after tamoxifen injection in βAdarHET and diabetic βAdarKO mice (n=3 for each genotype, 5 sections per mice 150mM apart analyzed). * p<0.05 and ** p<0.005; Student’s two-tailed t-test; error bars indicate mean ± SEM (standard error of the mean).

To study the effect of impaired RNA editing on beta-cell biology, we used mouse insulin promoter (Mip)-CreER; Adar^lox/lox^ mice to disrupt the Adar gene specifically in beta-cells of postnatal mice. To trace the fate of recombined cells we used a R26R^YFP^ reporter. We injected tamoxifen to 1-month-old MipCreER;R26R^YFP^, MipCreER R26R^YFP^;Adar^lox/+^ and MipCreER;R26R^YFP^;Adar^lox/lox^ mice to produce animals with beta-cell-specific Adar deficiency (βAdarKO), Adar heterozygosity (βAdarHET), or intact Adar (**Fig. 1C**). One week after tamoxifen injection we isolated islets, sorted YFP+ cells and performed RT-qPCR to assess Adar expression. Beta-cells from βAdarKO mice showed near-complete loss of the Adar floxed exon expression, whereas beta-cells from βAdarHET mice showed intermediate expression levels compared to wild type (**Fig. 1D**). We next performed RNA-sequencing and observed a significant reduction in RNA editing in βAdarKO beta-cells compared with βAdarHET. Interestingly, Adar heterozygosity resulted in reduced RNA editing compared with wild type, indicating that a full complement of Adar is needed to properly edit endogenous dsRNA structures (**Fig. 1D**).

Within two weeks of tamoxifen injection, 44% of βAdarKO mice (23/52) developed diabetes, whereas heterozygotes remained normoglycemic (**Fig. 1E, S1A)**. Similarly, intraperitoneal glucose tolerance test (IPGTT) revealed impaired glucose clearance in βAdarKO mice but not in βAdarHET mice (**Fig. S1B**). There was a gender bias in the diabetic phenotype with 70% of males as opposed to just 16% females showing glycemic deterioration – a phenomenon familiar from other settings of beta-cell damage, likely related to the protective effects of estrogen on female mouse islets (*26*) (**Fig. S1C and S2D**). The insulin-stained area in the pancreas of diabetic mice was greatly reduced (on average, by 87%), as was pancreatic insulin content (**Fig. S1E**), suggesting that in mutant mice, beta-cells were eliminated (**Fig. 1F**). Interestingly, the YFP-positive area was reduced to a lower extent (by 58% on average) (**Fig. 1F**), raising the possibility that some βAdarKO beta-cells have lost insulin (see below).

RNA editing has been shown in other systems to be crucial for suppression of innate immune system activation by dsRNA-binding cytosolic proteins (*4, 7*). To investigate whether beta-cell failure in βAdarKO mice was associated with islet inflammation, we stained pancreatic sections for the pan-leukocyte marker CD45 and scored islet infiltrates. Seven days after tamoxifen injection, 12.5% of islets in βAdarKO mice showed moderate to severe infiltration, and this score increased to 60% by 12-14 days. No insulitis was observed in βAdarHET mice (**Fig. 2A** and **Fig. S2A**). There were no gender-specific differences in the extent of islet infiltration (**Fig. S2B**). Thus, the greater incidence of diabetes in males cannot be accounted for by more severe islet inflammation. Islet infiltrates contained numerous Iba+ macrophages, with peri- and intra-islet presence of T cells (CD3+ cells, as well as some CD8+ cells), and fewer B cells (CD19+) (**Fig. 2B**). The islet infiltrate score correlated with glycemic deterioration in βAdarKO mice (**Fig. S2C**), suggesting that islet inflammation was responsible for beta-cell dysfunction and diabetes. To understand the molecular basis for islet inflammation, we performed RNA-sequencing on FACS-sorted YFP+ cells from βAdarKO and from βAdarHET mice, 7 days after tamoxifen injection. Adar deficiency induced a classic interferon response, with 69% of the 238 genes upregulated (FDR<0.05) known to be regulated by IFN (Interferome database (*27*)), including type I IFN genes (Ifna4, Ifnb1), ISGs such as Isg15 and Irf7, dsRNA sensors (Ifih1, Ddx58, Oas2 and Eif2ak2 [Pkr]) and the pro-inflammatory genes Ccl5 and Cxcl10 (**Fig. 2C**). Accordingly, the most significantly enriched gene sets among genes induced in βAdarKO beta-cells were alpha and gamma interferon pathways (**Fig. 2D and Fig. S2D**). In addition to the interferon-related pathways, gene sets including Myc targets, oxidative phosphorylation, TNFα pathway, and the unfolded protein response were significantly activated in beta-cells following Adar inactivation (**Fig. 2D**). Thus, Adar-deficient mouse beta-cells display an IFN response, including transcriptional activation of pro-inflammatory cytokines and chemokines that drive islet inflammation.

**Fig. 2:**
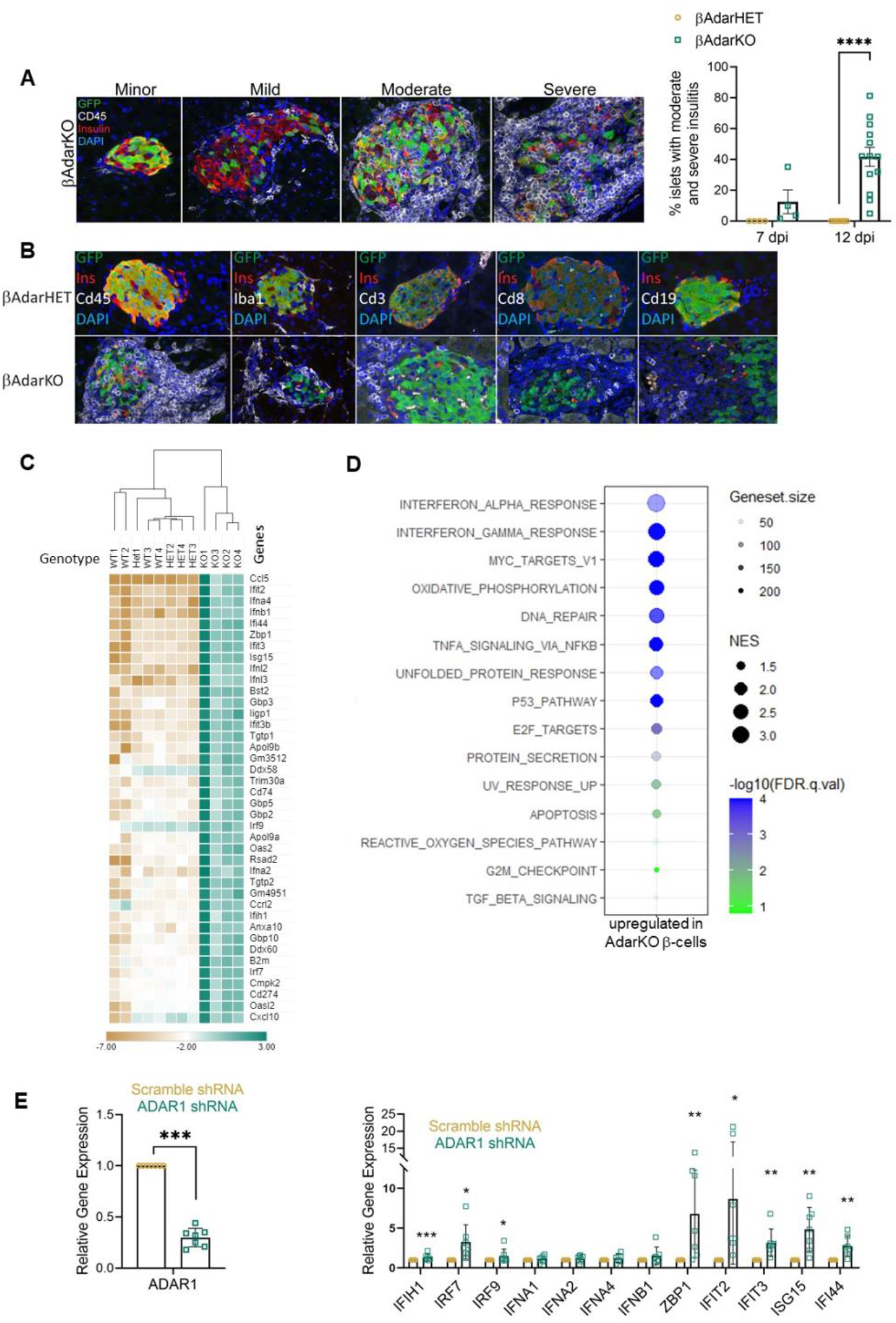
Insulitis and interferon response in βAdarKO mice. (**A**) Classification of islets from βAdarKO mice according to the extent of inflammatory infiltrates as assessed by immunostaining for insulin (β-cells), GFP (mutant β-cells) and CD4*5 (immune cel*l**s)**. Islets were scored 7 days and 12 days after tamoxifen injection. At least 100 islets from 5 different slides (80μm apart) were scored for each mouse. Each dot represents score for a single mouse. (**B**) Pancreas sections from βAdarHET (top) and βAdarKO (bottom) were stained for CD45 (pan-leukocyte marker), Iba1 (macrophages), CD3 (T cells), CD8 (T cells) and CD19 (B cells). (**C**) RNA-Seq heatmap representing ISG expression levels (after logarithmic transformation and row-centering) and hierarchical clustering (using Euclidean distance) in sorted β-cells from βAdarWT (WT), βAdarHET (HET) and βAdarKO (KO) mice. Brown and green reflect low and high expression levels, respectively, as indicated in the log2-transformed scale. (**D**) Bubble plot based on GSEA analysis showing top upregulated MSigDB hallmark gene sets in AdarKO compared to AdarHET β-cells. The bubble color, size and transparency represent -log10 (FDR), normalized enrichment score (NES) and gene set size, respectively. (**E**) ADAR1 knockdown in human islets induces ISG expression. RNA isolated from human pseudoislets (n=7 donors) transduced with either ADAR1 shRNA or scramble shRNA pseudoislets was assessed for expression of ADAR1 and interferon-stimulated genes by RT-PCR, with normalization to scramble shRNA samples. * p<0.05; ** p<0.01; *** p<0.001.

To examine the role of ADAR1 in human islets, we used a pseudoislet approach, where human islets are dispersed into singe cells, transduced with a shRNA ADAR1, and allowed to reaggregate over six days before analysis (**Fig. S3**). Knockdown of ADAR1 in human islets triggered an IFN response as demonstrated by the induction of several ISGs (**Fig. 2E**).

Interestingly, despite the reduction in Adar expression and RNA editing levels in βAdarHET mice compared with wild-type (**Fig. 1D**), Adar heterozygosity did not elicit a transcriptional IFN signature in beta-cells (**Fig. 2C**). These findings are consistent with normoglycemia and lack of islet inflammation in heterozygous mice and suggest that >50% reduction in Adar activity is needed to cross a threshold – presumably amount of dsRNA – for triggering an interferon response and consequently inflammation in mouse beta-cells.

Taken together, Adar deficiency in mouse or human beta-cells triggers a massive interferon response, which *in vivo* is associated with islet infiltration by immune cells, beta-cell dysfunction and loss, and diabetes.

To better characterize the phenotype of beta-cells following Adar disruption, we performed immunofluorescence staining on pancreas sections from βAdarHET and normoglycemic βAdarKO mice 12-14 days after tamoxifen injection. Non-diabetic mice were chosen to avoid potentially confounding effects of hyperglycemia on beta-cell gene expression. We co-stained sections for CD45, the GFP tracer and key beta-cell specific markers. We also took advantage of the incomplete activation of the CreER transgene in beta-cells to analyze the effects of Adar disruption in mosaic islets at a single cell resolution. This strategy can help distinguish paracrine effects (e.g. mediated by signals from inflammatory or cells) from cell-autonomous responses (caused intrinsically by Adar mutation).

As shown in **Fig. 3A-C**, insulin staining intensity per beta-cell was markedly reduced in βAdarKO mice, though strikingly this was exclusive to inflamed islets (i.e., non-inflamed islets appeared normal, even if they contained a majority of Adar-deficient beta-cells as evident from GFP staining). Similarly, the levels (per cell) of the insulin-processing enzyme PC1/3 were lower in inflamed mutant islets, compared with non-inflamed islets in βAdarKO or βAdarHET mice (**Fig. 3D**). Importantly, this phenotype was seen in both Adar-deficient beta-cells (marked as GFP^+^) and Adar-proficient beta-cells (GFP^-^). In contrast, protein levels of key beta-cell transcription factors Pdx1, Nkx6.1 and Pax6 in infiltrated islets were normal (**Fig. S4A-H**).

**Fig. 3:**
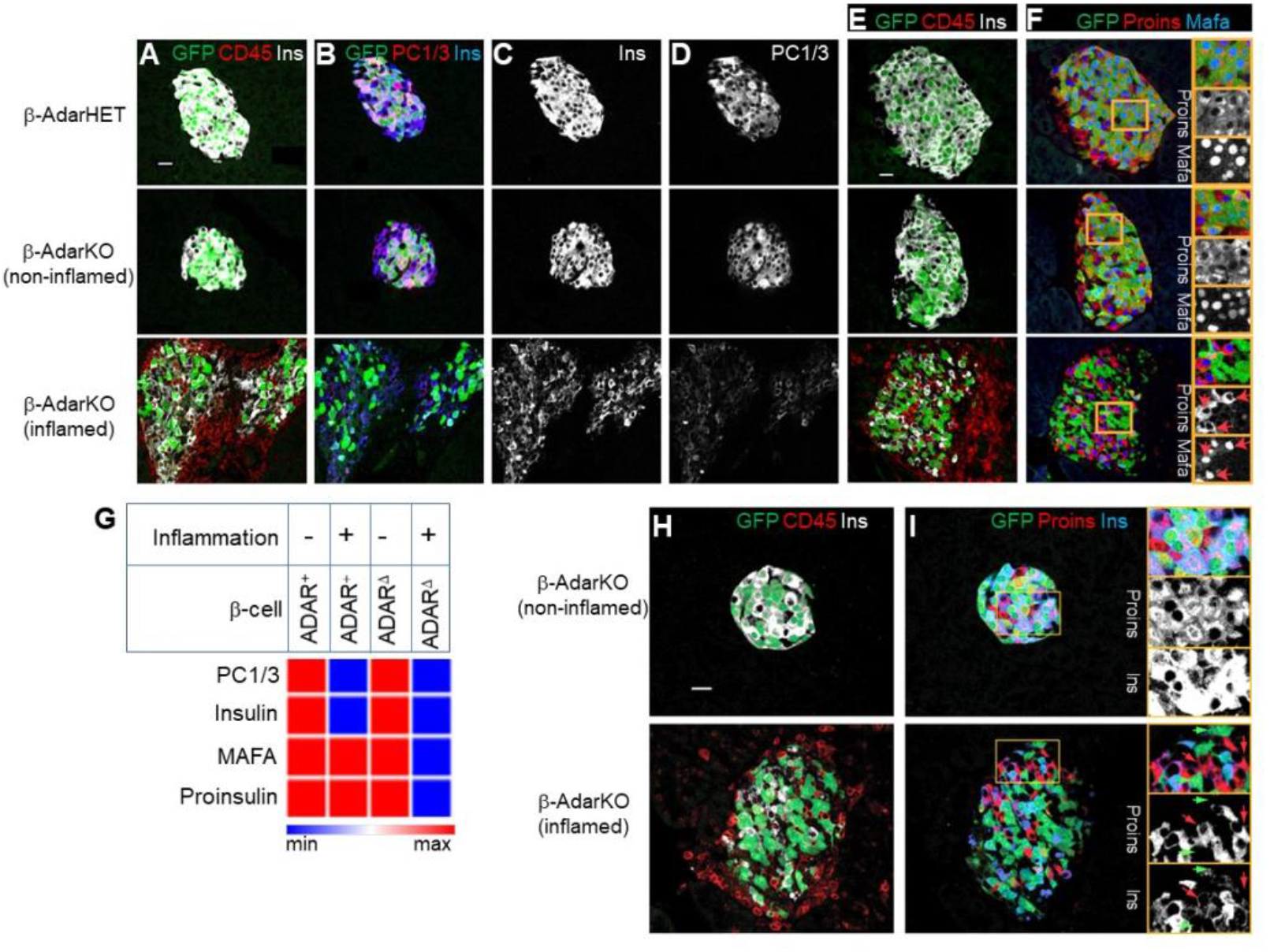
Adar inactivation combined with islet inflammation alters beta-cell expression program. Pancreatic sections from non-diabetic βAdarKO and control βAdarHET mice, 12-14 days after Tamoxifen injection were immunostained for GFP, CD45 and beta-cell markers. (**A-D**) Pancreatic sections from βAdarHET (upper panel) and βAdarKO mice (mid and lower panel) were immunostained for GFP, CD45 and Insulin (**A**) and serial sections stained for GFP, Insulin and PC1/3 (**B-D**). Representative micrographs of islets demonstrate reduced insulin and PC1/3 staining in inflamed islets of βAdarKO mice (**C-D**, lower panels). (**E, F**) Pancreatic sections from βAdarHET (upper panel) and βAdarKO mice (mid and lower panel) were immunostained for GFP, CD45 and Insulin (**E**) and serial sections stained for GFP, proinsulin and MafA (**F**). Insets show dramatic reduction in MafA coinciding with reduced proinsulin levels in Adar-deficient (GFP-positive) beta-cells of inflamed islets, but not in their neighboring Adar-positive GFP-negative counterparts (marked by red arrows) (**F**, lower panel). (**G**) Scheme depicting how Adar inactivation and inflammation affect beta-cell marker expression. (**H, I**) Pancreatic sections from βAdarKO mice were immunostained for GFP, Insulin and CD45 (**H**) and serial sections were stained for GFP, proinsulin and insulin (**I**). Representative micrographs from non-inflamed (**H**) and inflamed (**I**) islets from the same section show that in inflamed islets, Adar-knockout GFP-positive beta-cells (marked by green arrows) exhibit lower levels of insulin and proinsulin, while Adar-positive GFP-negative beta-cells (red arrows) retain high levels of proinsulin together with reduced insulin levels. Scale bars, 20 micrometers.

Thorough examination of infiltrated islets showed that whereas levels of the MafA transcription factor were dramatically reduced in mutant (GFP^+^) beta-cells, they were preserved in adjacent wild-type beta-cells (**Fig. 3E-F**). This bimodal pattern of expression coincided with proinsulin expression: within inflamed islets of βAdarKO mice, GFP^+^ (mutant) beta-cells were typically devoid of proinsulin, while YFP^-^ (wild-type) beta-cells retained high levels of proinsulin (**Fig. 3E-F**).

Thus, Adar deficiency in beta-cells exerts two distinct phenotypes related to insulin expression, which are both associated with islet inflammation (**Fig. 3G**). In Adar-deficient beta-cells, the levels of MafA and proinsulin are reduced. Since MafA is a key transcription factor of insulin, its downregulation is a likely reason for the reduction in proinsulin. In both Adar-deficient and wild-type beta-cells, the levels of PC1/3 and mature insulin are reduced. Since PC1/3 is essential for the processing of proinsulin to mature insulin, its downregulation is a potential cause for the reduction in insulin. We note that the loss of insulin was seen as early as 7 days after tamoxifen injection; given the long half-life of insulin (around 3 weeks (*28*)), this suggests that degradation of pre-existing mature insulin was accelerated in inflamed islets of these normoglycemic mice, regardless of autonomous Adar deficiency.

In summary, our results indicate that reduced levels of MafA and proinsulin in mutant beta-cells are caused by a combination of cell autonomous determinants (driven by Adar deficiency) and autocrine/paracrine signals (triggered by IFN produced by Adar-deficient beta-cells or additional cytokines originating from inflammatory cells). In contrast, reduced levels of PC1/3 and mature insulin are caused purely by paracrine signals, irrespective of Adar status.

The interplay between paracrine/autocrine signals and cell autonomous cues results in the cohabitation within the same inflamed islets of proinsulin-deficient, insulin-deficient Adar-negative beta-cells, together with Adar-positive beta-cells that exhibit high proinsulin and dramatically reduced insulin levels (**Fig. 3H-I**). As elaborated in the Discussion, these phenotypes bear striking resemblance to the situation in islets from patients with recent onset T1D, where proinsulin^+^ insulin^-^ beta-cells (“empty beta-cells”) are often observed (*29, 30*). Strikingly and in contrast to the impact of inflammation on beta-cell phenotype, alpha- and delta-cells within inflamed islets of mutant mice were largely spared (**Fig. S4I-L**) consistent with the observed phenotype in human T1D.

Our *in-vivo* observations cannot determine if inflammation is causative or is merely a biomarker of islets producing particularly high levels of type I interferon, which is both fueling inflammation and causing MafA and proinsulin loss in mutant beta-cells (and possibly PC1/3 and insulin loss in all beta-cells). To better dissect the signaling pathways underlying misexpression of beta-cell markers in our model, we performed *ex-vivo* experiments on islets from βAdarKO and βAdarHET mice, isolated 7 days after tamoxifen injection. Culturing βAdarKO islets *ex-vivo* for 3 additional days caused reduced MafA and proinsulin levels in AdarKO GFP^+^ β-cells, thus recapitulating the phenotype observed in inflamed islets *in-vivo* (**Fig. 4A-B)**. In contrast, the levels of insulin protein were not significantly altered in GFP^+^ β-cells (**Fig. 4B)**. Strikingly, treatment with an inhibitor of the Janus kinase/signal transducer and activator of transduction (JAK/STAT) signaling pathway (ruxolitinib) maintained normal levels of MafA and proinsulin in AdarKO β-cells, demonstrating that this phenotype is dependent on IFN signaling (**Fig. 4A-B)**. These results indicate that inflammatory cells are not required for the cell-autonomous phenotype observed in Adar-deficient beta-cells. Rather, we propose that MafA and proinsulin are reduced due to a combination of autonomous Adar deficiency and autocrine or paracrine IFN signaling. Finally, Adar knockdown in human islets led to reduced RNA levels of Insulin, MafA, MafB, Pdx1 and Nkkx6.1 (**Fig. 4C**), and reduced MafA protein levels (**Fig. S4M**), indicating that inflammatory infiltrates are not essential for altering beta-cell identity genes in human islets.

**Fig. 4:**
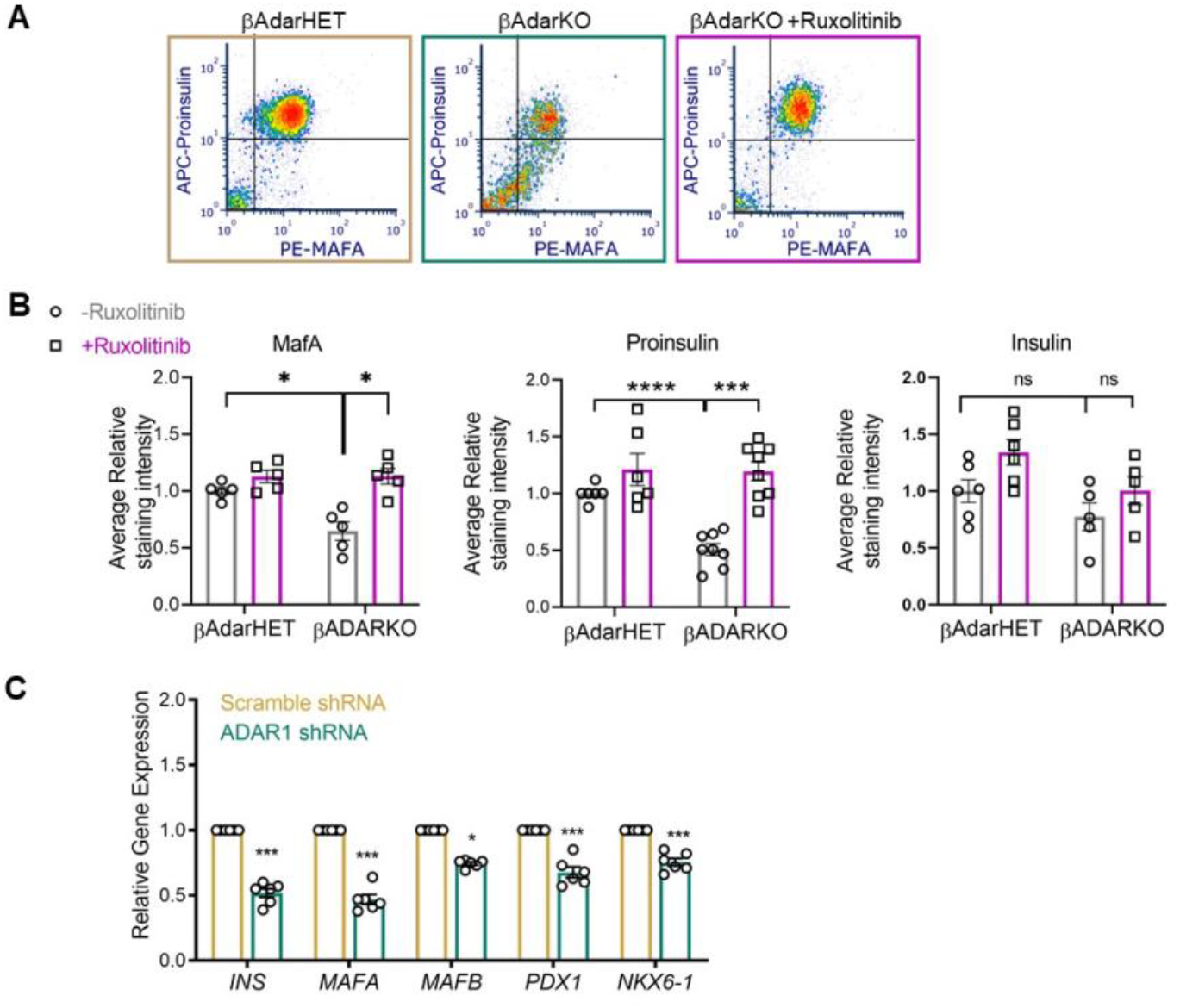
Adar inactivation causes decreased expression of beta-cell markers in mouse and human islets ex*-*vivo. (**A-B**) βAdarHET and βAdarKO islets were isolated 7 days after Tamoxifen injection and cultured ex-vivo for three additional days with or without ruxolitinib (4μM), a JAK/STAT inhibitor before immunostaining for insulin, proinsulin and GFP and FACS analysis. (**A**) Representative FACS dot plot of GFP-positive beta-cells reveals decrease in proinsulin and MafA levels in AdarKO beta-cells compared to AdarHET beta-cells. Ruxolitinib prevented the reduction in proinsulin and MafA levels in GFP-positive AdarKO β-cells. (**B**) Quantification of average MafA, proinsulin and insulin staining levels in GFP-positive beta-cells from 5-6 independent experiments shows significant reduction in MafA and proinsulin levels but not in insulin levels in AdarKO compared to AdarHET β-cells and in untreated versus ruxolitinib treated AdarKO GFP-positive beta-cells. *p<0.05, ***p<0.001, ****p<0.0001, ns: non-significant; Student’s two-tailed t-test; error bars indicate mean ± SEM. (**C**) Expression of islet cell-enriched transcription factors is reduced in ADAR1 knock down human islets (n=7 donors). * p<0.05, *** p<0.01.

The onset of insulin treatment in T1D is often associated with a temporary remission (the honeymoon period) (*31*). Similarly, insulin therapy (reducing glycemic load and the drive for secretion) or treatment with diazoxide (a KATP channel opener that prevents glucose-induced membrane depolarization and insulin secretion) in rodent models of T1D have been reported to reduce the incidence of diabetes (*32, 33*). These findings gave rise to the concept of “beta-cell rest” – a beneficial effect of reduced beta-cell workload. This phenomenon is typically interpreted in the context of reduced lipo- and glucotoxicity, which improves beta-cell survival and function (*34, 35*).

We hypothesized that reduced beta-cell workload, and consequently reduced glucose-dependent signaling (**Fig. 5A**), may also affect the interferon response of beta-cells, and hence directly impact their interaction with the immune system. To test this idea, we isolated islets from βAdarKO mice three days after tamoxifen injection (prior to the onset of islet inflammation), cultured the islets in different glucose concentrations for three additional days, and measured the expression of selected ISGs using qRT-PCR. Culturing βAdarKO islets in 11mM glucose led to a dramatic elevation of ISGs compared with a culture in 5mM (**Fig. 5B**), while βAdarHET islets did not show ISG expression in either concentration of glucose (**Fig. S5**). We also treated βAdarKO islets with a small molecule glucokinase activator (GKA), a drug that increases the affinity of glucose to glucokinase thereby increasing the rate of glycolysis (*36*). GKA led to elevated expression of ISGs even in the presence of 5mM glucose (**Fig. 5B**), indicating that the rate of glycolysis rather than glucose itself is driving ISG expression in βAdarKO islets.

**Fig. 5:**
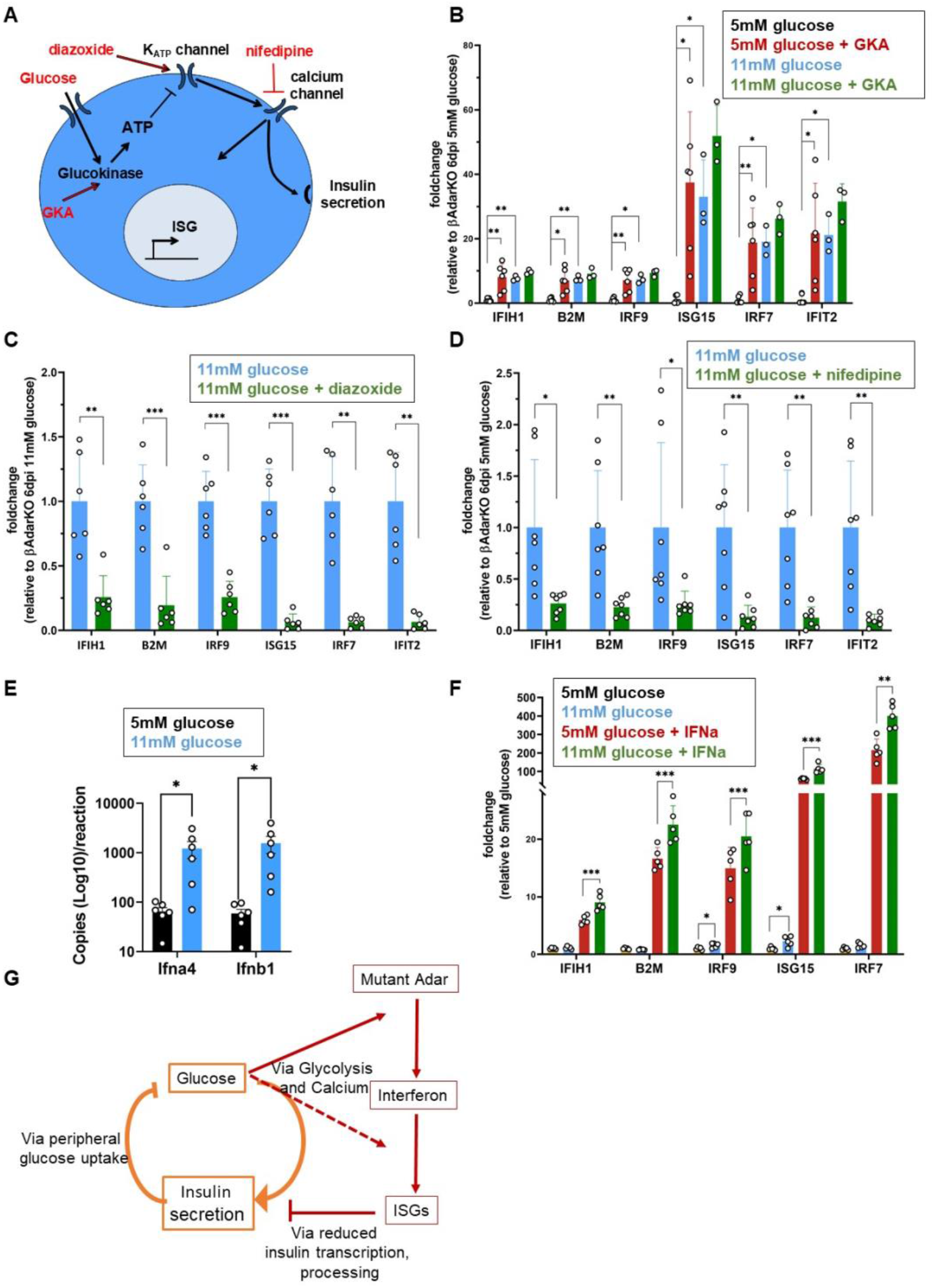
Glucose metabolism and calcium influx enhance the interferon response in islets from βAdarKO mice. **(A)** Graphical model depicting glucose metabolism and glucose-induced pathways to insulin secretion. Pharmacological interventions that modulate insulin secretion (marked in red) were tested to delineate necessary pathways for glucose-induced ISG expression in βAdarKO islets **(B-D)** Islets from βAdarKO mice were isolated 3 days after tamoxifen injection and cultured for three days in RPMI medium supplemented with 5mM or 11mM glucose and pharmacological modulators of insulin secretion, as indicated (10uM GKA treatment (B), 325uM diazoxide (C), and 10uM nifedipine (D). ISG mRNA expression was assayed by RT-qPCR and normalized to Actb gene expression. **(E)** Glucose activates transcription of type I IFN genes in islets from βAdarKO mice. Islets from βAdarKO mice were isolated 3 days after tamoxifen injection and cultured for three days in RPMI medium supplemented with 5mM or 11mM glucose. Expression of *Ifna4* a*nd Ifnb1 genes w*as assessed by RT-ddPCR (Droplet Digital PCR). *p<0.05; Student’s paired t-test; error bars indicate mean ± SEM. **(F)** Glucose modulates the interferon response in wild-type islets. Islets from βAdarWT mice were cultured for three days in 5mM or 11mM glucose and treated for 24 hours with 800u/ml IFNα. ISG expression was assayed by RT-qPCR and normalized to Actb expression. n>=6, *p<0.05, **p<0.01, ***p<0.001; Student’s unpaired two-tailed t-test; error bars indicate mean ± SEM. **(G)** Model illustrating the putative vicious cycle of interferon response and hyperglycemia, via beta-cell workload that amplifies the interferon effect.

To delineate the signaling pathway that regulates the interferon response in βAdarKO islets downstream of glycolysis, we treated βAdarKO islets with pharmacological activators or inhibitors of steps in the classical glucose-stimulated insulin secretion pathway (**Fig. 5A**). Diazoxide prevented the induction of ISGs by high glucose (**Fig. 5C**). Downstream of membrane depolarization, the calcium channel blocker nifedipine blocked ISG induction by high glucose (**Fig. 5D**).

Together our results indicate that glucose-induced calcium influx is necessary for ISG expression triggered by deficient RNA editing.

We performed additional experiments to better characterize the molecular mechanism underlying the synergistic effect of Adar inactivation and high glucose on the beta-cell interferon response. The interferon response to dsRNA has cell-autonomous and autocrine/paracrine phases. Initially, dsRNA accumulates and activates dsRNA sensors including IFIH1 which initiate a signaling pathway culminating in IRF3-mediated transcription of type I interferon (IFN-α/β) genes. Interferon is then secreted and sets off a second phase of autocrine/paracrine signaling events, which stimulate the expression of ISGs via the interferon receptor and JAK/STAT pathway (*2, 37*).

To investigate whether the first, cell-autonomous phase is impacted by glucose, we measured *Ifna4* and *Ifnb1* transcripts (encoding IFN-α/β) in βAdarKO islets cultured in 5mM or 11mM glucose. Increasing glucose concentration augmented *Ifna4* and *Ifnb1* mRNA levels by 26 and 38-fold, respectively (**Fig. 5E**). To test the effect of glucose on the paracrine phase, we treated wild-type islets with interferon-α and cultured the islets in different glucose concentrations. High glucose (11 mM) boosted the ISG expression in response to IFN treatment about 2-fold, revealing that it enhances JAK/STAT signaling independently of RNA editing status (**Fig. 5F**). We conclude that glucose flux mainly regulates the cell-autonomous signaling cascade from Adar deficiency to induction of type I IFN in beta-cells, and has a minor but significant effect on the response of beta-cells to exogenous interferon.

In summary, glucose metabolism through calcium signaling is a strong amplifier of the interferon response in beta-cells, suggesting a novel link between metabolism and islet inflammation.

## Discussion

Our findings reveal that Adar activity in beta-cells is an essential safeguard against the development of aberrant interferon response and islet inflammation. We found that reduced ADAR in mouse and human islets leads to interferon-stimulated gene expression. In mice lacking ADAR in beta-cells, islets are targeted by inflammatory cells, resembling insulitis in human type 1 diabetes. These islet-related phenotypes are consistent with the inferred functions of ADAR in other tissues, and likely result from IFIH1 sensing of unedited endogenous dsRNA species. The evolutionary logic is that Adar-mediated dismantling of endogenous dsRNA (which have accumulated historically in the genome via retroelement insertions in expressed genes) allows cells to discriminate between self and exogenous viral dsRNA and selectively activate innate immunity in response to viral infection (*4, 7*). Beyond these universal roles of Adar, our results show that the combination of ADAR deficiency and inflammation dramatically impairs the molecular identity of beta-cells (both βAdarKO and adjacent wild-type beta-cells). Adar-deficient beta-cells lose expression of MafA, and, likely as a consequence, have reduced levels of proinsulin. We observed these phenotypes *in vivo* only in inflamed islets. Adar deficiency *in vitro* led to reduced MafA and proinsulin expression through JAK/STAT signaling, revealing necessity of IFN rather than inflammation (potentially reflecting islets with high levels of dsRNA).

Adar deficiency also had dramatic effects on wild-type beta-cells within inflamed islets. While these cells retained beta-cell identity, they lost expression of PC1/3 and mature insulin – the latter likely resulting from deficient proinsulin processing and possibly accelerated insulin degradation. Similar phenotypes resulting from the action of pro-inflammatory cytokines have been noted in T1D (*29, 38, 39*).

Overall, the combination of autonomous and non-autonomous phenotypes following ADAR disruption in beta-cells bears a striking similarity to phenotypes observed in the pancreas of humans with T1D (see below).

Our discovery that the beta-cell interferon response is influenced by glucose metabolism potentially explains the association between beta-cell workload and the progression of early stage T1D. The lowering of glycemic load clearly has a direct impact on glucotoxicity via reduced ER or oxidative stress (*40–42*). However, it has been difficult to examine if immune responses are also affected. Focusing on the transcriptional interferon response of beta-cells, without involvement of either innate or adaptive immune cells, allowed us to observe that glucose metabolism and downstream steps in the classical insulin secretion pathway are key modulators of the beta-cell response to deficient RNA editing, in at least two distinct steps. First, glucose enhances the interferon response to loss of Adar. This can occur via the massive transcriptional response of beta-cells to glucose, which could lead to elevated expression of genes with particularly “dangerous” dsRNA configurations when unedited. Alternatively, glucose may enhance the signaling pathway from dsRNA sensing to the primary interferon response. Secondly, glucose enhances moderately but significantly the transcriptional response of beta-cells to exogenous interferon, even in wild-type beta-cells that do not have excessive dsRNA. While the molecular mechanism mediating this effect remains to be elucidated, it suggests an interesting positive feedback loop acting in a particularly sensitive stage during the development of T1D. Interferon and other cytokines can reduce beta-cell functionality (e.g. via reduced levels of PC1/3 and mature insulin, as shown here); this in turn would render systemic glucose clearance less efficient, and increase the workload on beta-cells which will drive a stronger interferon response (model, **Fig. 5G**). Note that the process can take place while the individual is still euglycemic, potentially progressing silently until erupting when functional beta-cell mass can no longer sustain euglycemia. This description is consistent with the natural phenomenon of the honeymoon period in recent-onset T1D, with pre-clinical and clinical reports on the benefits of enforced beta-cell rest, and with a recent report that the calcium channel blocker verapamil improves beta-cell function in patients with recent-onset T1D (*43*). The vicious cycle of inflammation and increased workload also suggests targets for early intervention. For example, low dose insulin has been examined as a preventive therapy for T1D, mainly aimed at inducing immunotolerance (*44, 45*). The mechanism proposed here supports the concept seen in the Diabetes Control and Complication Trial that higher doses of insulin preserve glucose-stimulated insulin secretion (*46*), suggesting that insulin treatment would reduce beta-cell metabolism and the islet interferon response in people at risk to develop T1D.

A recent study has mapped human genetic variants that are associated with reduced editing of endogenous dsRNAs (RNA editing Quantitative Trait Loci, or edQTLs), and linked these variants to the risk of developing autoinflammatory and autoimmune disease, including T1D (*18*). This suggests that endogenous, rather than virus-derived dsRNA molecules, are important upstream activators of the interferon response leading to T1D (*47, 48*). However, the cell type(s) mediating the effect and the molecular mechanisms leading to beta cell damage are not known. Our study used Adar disruption as a model system to reliably induce dsRNA-mediated interferon response in beta-cells *in vivo* and to examine its downstream effects. Unexpectedly, the consequences of Adar deficiency in beta-cells *in vivo* resemble features of early stage T1D, which are typically explained as consequences of antigen-specific autoimmunity. First, although Adar deficiency brought about massive islet inflammation, only beta-cells within islets were destroyed. This phenotype does not result simply from the autonomous requirement for Adar in beta-cells, since adjacent wild-type beta-cells were also affected. Selective destruction of beta-cells is a hallmark of T1D (*29, 48*). While antigen-specific autoreactive T cells are clearly key players in T1D, our findings suggest that beta-cell selectivity can emerge also in the absence of autoreactive lymphocytes, presumably via vulnerability brought about by distinct metabolic wiring of beta-cells. Second, islets with mosaic deletion of Adar in beta-cells had abundant empty, or “ghost” beta-cells, which contained proinsulin but lacked mature insulin. Such cells are observed in the pancreas of patients with recent-onset T1D (*29, 39*). Moreover, in both βAdarKO mice and human T1D, lack of mature insulin is associated with reduced levels of the PC1/3 convertase (*49*). Our study suggests that PC1/3 is reduced due to exposure to paracrine signals emanating from either mutant beta-cells or infiltrating immune cells. Third, we observed a highly heterogenous inflammatory response, with some islets being heavily infiltrated and some completely spared, despite efficient Adar deletion. Mosaic islet infiltration is an intriguing, unexplained phenomenon in T1D (*30, 50*). We propose that it results from diverging outcomes of an early interferon response in a subset of beta-cells, which need to cross a certain threshold of intensity to attract inflammatory cells. This in turn may exacerbate local islet inflammation. The possibility that islets differ in their metabolic activity may further drive heterogenous islet inflammation (*51, 52*).

In summary, we describe the dependency of mouse and human beta-cells on intact Adar function. In the absence of Adar, editing of endogenous mRNA fails, leading to the accumulation of dsRNA, activation of an interferon response, islet inflammation and diabetes. The consequences of Adar1 deficiency in beta-cells resemble key aspects of early T1D, and are consistent with recent genetic evidence that implicates RNA editing in human inflammatory diseases including autoimmune diabetes. Together these findings suggest a model for T1D in which defective RNA editing of beta-cell transcripts leads to an anti-viral response and destructive islet inflammation (**Fig. S6)**. Finally, the metabolic dependency of the interferon response suggests the presence of a positive feedback loop that may drive progression of T1D, with potential implications for pharmacologic intervention.

## Acknowledgments

This study used human pancreatic islets that were provided by the NIDDK-funded Integrated Islet Distribution Program at the City of Hope (DK098085).

## Author contributions

Conceptualization, YD, AK, EYL, CD, ACP;

Investigation, EK, SP, AK, CD, RCF;

Writing and Editing, AK, YD, BG, EYL, CD, ACP;

Review & Editing, all authors;

Supervision, YD, AK, EYL, ACP; Funding, YD, ACP, EYL.

## Data and materials availability

The RNA-seq data is available at XXX._This paper does not report original code. Any additional information required to reanalyze the data reported in this paper is available from the lead contact upon request.

## Supplementary Materials

Materials and Methods

Figs. S1 to S7

Table S1

References (51-59)

## Supplementary Materials

### Materials and Methods

#### Mouse Models

All experiments presented in this study were conducted according to regulatory standards approved by the joint ethics committee (IACUC) of the Hebrew University and Hadassah Medical Center. The Hebrew University is an AAALAC International-accredited institute. The mouse strains used in this study were ROSA26-LSL-EYFP (Jackson Laboratory), MipCreER (*51*) and Adar^lox^ mice (with loxP-flanked Adar) (*8*). Tamoxifen (20 mg/ml in corn oil; Sigma-Aldrich) was injected subcutaneously into 1-month-old adult mice. Two doses of 8mg (2 days apart) were used to achieve maximal recombination in beta cells. For Intraperitoneal Glucose Tolerance Test (IP-GTT), mice were fasted overnight and glucose (2g/kg) was injected intraperitoneally. YFP+ cells were strictly insulin+ in βAdarHET mice, verifying beta cell specificity of the MIP-CreER driver (**Fig. S7**). The average rate of recombination in beta cells was 67% (data not shown).

#### Human islets and pseudoislets

Human islets from IIDP (https://iidp.coh.org) were used to create pseudoislets that were handled and assessed for gene or protein expression and hormone secretion by perifusion as previously described (*52*). Donor information is provided in **Table S1**. Briefly, hand-picked human islets (>95% purity) were cultured in CMRL 1066 medium and dispersed with HyClone trypsin (Thermo Scientific). The dispersed islet cells were incubated either with CMV-mCherry scramble control vector adenovirus or CMV-mCherry-U6-ADAR1vector virus at a multiplicity of infection of 500 for 2 hours in Vanderbilt Pseudoislet Media and then washed and plated in Cell-Carrier Spheroid Ultra-low attachment microplates (PerkinElmer) with 2000 cells per well. After 6 days of culture, pseudoislets were harvested for subsequent analysis. We confirmed that viral transduction of human islet cells did not induce an interferon response (**Fig. S8)**.

#### Immunostaining

Paraffin sections (5-μm-thick) were prepared from formalin-fixed, paraffin-embedded pancreata. Sections after rehydration and antigen retrieval were immunostained overnight at 4°C with primary antibodies (see key resources table) in CAS-block blocking solution (Thermo fisher). For fluorescent immunostaining, slides were stained with Cy™2, Cy™3 and Cy™5-conjugated AffinityPure Donkey IgG (H+L) secondary antibodies (Jackson ImmunoResearch). DNA was counterstained using DAPI (Sigma-Aldrich). Immunofluorescence images were captured using a Nikon C2 or Olympus FV1000 confocal microscope. To determine beta cell area, consecutive paraffin sections 75 μm apart spanning the entire pancreas (approximately 9 sections/pancreas) were stained for insulin or YFP (using an anti-GFP antibody) and hematoxylin (SIGMA); slides were incubated with Biotin-SP-conjugated AffinityPure Donkey IgG (H+L) (Jackson ImmunoResearch), followed by Avidin HRP (Sigma-Aldrich) and 3,3′-Diaminobenzidine (DAB) substrate (Thermo Scientific). Digital images of sections at a magnification of ×40 were obtained and stitched using Nikon-TL and NIS-Elements software.The fraction of tissue covered by insulin or YFP staining was determined using ImageJ software.

#### Insulitis

Tissue sections (5 μm) were immunostained for YFP (using an anti-GFP antibody), insulin and CD45. Images of islets from five noncontiguous sections of each pancreas were captured on a Nikon C2 and islets scored for infiltrating leukocytes as shown in **Fig. 2A**: no infiltration, minor (less than 10 peri-islet leukocytes present at any point around islet), mild (>30% peri-islet infiltrate), moderate (intra-islet infiltrate) or severe (>50% intra-islet infiltrate). At least 100 islets/mouse with n = 4–13 mice/genotype were scored and averaged to calculate insulitis score.

#### Pancreas Insulin Content

Pancreata from βAdarWT, βAdarHET or diabetic βAdarKO were harvested 16 days after tamoxifen injection and homogenized in ice-cold acidic alcohol (0.18N Hydrochloric Acid (HCl), 70% Ethanol). After 24 hours incubation at 4°C tissue homogenates were centrifuged, supernatant insulin concentrations were measured by an enzyme-linked immunosorbent assay (ELISA) (Crystal Chem Inc.) and normalized to tissue weight.

#### Mouse Islets

Mouse islet isolation was performed as previously described (*53*). Briefly, islets were isolated using collagenase P (Roche) injected into the pancreatic duct, followed by Histopaque gradient (1119 and 1077; Sigma-Aldrich). For ex-vivo assays, 30–40 handpicked islets per replicate were incubated in RPMI-1640 medium (Biological Industries) supplemented with 5mM or 11mM glucose, 10% FBS (Sigma-Aldrich), L-glutamine and penicillin-streptomycin in a 37 °C, 5% CO2 incubator and treated for the indicated times with 10μmol/L Glucokinase activator (GKA, Pfizer), 325 μmol/L diazoxide (Sigma-Aldrich), 10 μmol/L nifedipine (Sigma-Aldrich) or 800 units/ml Interferon alpha (IFNa, BioLegends).

#### Flow Cytometry and Cell Sorting (FACS)

For live cell sorting, islets from βAdarWT, βAdarHET or βAdarKO mice were dissociated to single cells using Accumax (Sigma-Aldrich), stained with DAPI (0.2ug/ml) for dead cell discrimination, and DAPI-negative, YFP-positive cells sorted on a FACS Aria III (Becton Dickinson). For flow cytometry analysis, dissociated islet cells were fixed and permeabilized using Cytofix/Cytoperm and Perm/Wash (BD Biosciences). Cells were stained in Perm/Wash buffer with primary antibodies, followed by secondary antibodies (Cy2, Cy3 or Cy5 conjugated; Jackson ImmunoResearch). Cells were analyzed using an LSR-Fortessa-Analyzer with at least 10,000 events recorded. Analysis of the results was performed using FCS-express 7 (De Novo Software).

#### RNA Extraction

RNA was isolated using Zymo Direct-Zol™ RNA-MicroPrep kit and reverse transcription was performed with a qScript kit (Quanta Biosciences) according to the manufacturer’s instructions. Quantitative real-time PCR (qRT-PCR) was performed on a CFX96 Real-Time System (Bio-Rad) with PerfeCTa SYBR Green SuperMix (Quanta Biosciences) and gene-specific primers (see key resources table). RT-Droplet Digital PCR was performed on a ddPCR system (Bio-Rad) with Supermix, Droplet generation oil for probes (Bio-Rad) and gene-specific primers and probe (see key resources table).

Human pseudoislet RNA was extracted using Invitrogen RNAquous-Micro Total RNA Isolation kit (Thermo Fisher #AM1931) and quantified by Qubit Fluoremeter 2.0 as described. RNA integrity was confirmed (RIN 8.2-10, average 9.7±0.13) by 2100 Bioanalyzer (Agilent). cDNA was synthesized using high-capacity cDNA Reverse Transcription Kit (Applied Biosystems, 4368814) according to the manufacturer’s instructions. Quantitative PCR (qPCR) was performed using TaqMan assays and reagents from Applied Biosystems as described (*54, 55*).

Relative changes in mRNA expression were calculated by the comparative deltaCt method using Applied Biosystem StepOnePlus system.

#### RNA-Seq Library Preparation and Sequencing

Libraries were prepared using the SMARTer Stranded Total RNA-Seq Kit v3 - Pico Input Mammalian (Takara). Samples were sequenced (paired-end (2 × 75 bp)) on a NextSeq 500 (Illumina).

Raw FASTQ quality was assessed using FastQC (version 0.11.8), and barcodes were preprocessed using UMI-tools versions 1.1.1 and 1.1.2 (*56*). Mate 2 reads were further trimmed to 60 bases using seqtk trimfq with -b 7. Fastq files were then uniquely aligned to reference genome using STAR (version 2.7.3a) (*57*) with parameters --alignSJoverhangMin 8 -- alignIntronMax 1000000 --alignMatesGapMax 600000 --outFilterMismatchNoverReadLmax 1 -- outFilterMultimapNmax 1 and deduped using UMI-tools. Reads were counted with the htseq-count script version 2.0.1 (*58*) and normalized using DeSeq2 version 1.28.1 (*59*).

#### Detection and quantification of A-to-I RNA Editing

RNA Editing Index version 1.0 was used to assess the overall editing in *Alu* and *B1, B2* elements in human and mouse, respectively (*23*). This measure calculates the average editing level across all adenosines in repetitive elements weighted by their expression, thereby quantifying the ratio of A-to-G mismatches over the total number of nucleotides aligned to repeats and comprising a global, robust measure of A-to-I RNA editing.

#### Quantification and statistical analysis

Experimental values are presented as mean ± SEM. Unless stated otherwise, statistical significance was determined by two tailed unpaired Student’s t test. P < 0.05 was considered statistically significant (*p<0.05, **p<0.01, ***p<0.001, **** p<0.0001).

##### Antibodies and reagents

| REAGENT or RESOURCE | SOURCE | IDENTIFIER |
| --- | --- | --- |
| Antibodies |  |  |
| Guinea pig anti Insulin | DAKO | Cat# IR002 |
| Goat anti GFP | Abcam | Cat# Ab6673 |
| Rabbit anti Iba1 | Abcam | Cat# Ab178846 |
| Rat anti CD3 | Abcam | Cat# Ab11089 |
| Rat anti CD8 | Invitrogen | Cat# 14-0808-80 |
| Rat anti CD19 | Invitrogen | Cat# 14-0194-82 |
| Rabbit anti CD45 | Abcam | Cat# Ab10558 |
| Rabbit anti MafA | Bethyl | Cat# A700-067 |
| Mouse anti Proinsulin | Hybridoma bank | Cat# GS-9A8 |
| Rabbit anti PC1/PC3 | Abcam | Cat# Ab220363 |
| Mouse anti glucagon | Abcam | Cat #AB10988 |
| Rabbit anti mCherry | Abcam | Cat# AB167453 |
| Rabbit anti ADAR1 | Invitrogen | Cat # PA5-21369 |
| Rabbit anti somatostatin | Abcam | Cat# 108456 |
| Alexa Fluor® 488 AffiniPure Donkey Anti-Goat IgG (H+L) (min X Ck, GP, Sy Hms, Hrs, Hu, Ms, Rb, Rat Sr Prot) | Jackson ImmunoResearch Labs | Cat# 705-545-147, RRID:AB_2336933 |
| Cy3-AffiniPure Donkey Anti-Rabbit IgG (H+L) (min X Bov,Ck,Gt,GP,Sy Hms,Hrs,Hu,Ms,Rat,Shp Sr Prot) | Jackson ImmunoResearch Labs | Cat# 711-165-152, RRID:AB_2307443 |
| Alexa Fluor 647-AffiniPure Donkey Anti-Mouse IgM, $\mu$ Chain Specific (min X Hu,Bov,Hrs,Rat Sr Prot) antibody | Jackson ImmunoResearch Labs | Cat# 715-605-140, RRID:AB_2340861 |
| Chemicals, peptides, and recombinant proteins |  |  |
| Tamoxifen | Sigma-Aldrich | Cat# T5648-5G |
| PerfeCTa SYBR Green FastMix | Quanta Biosciences | Cat# 95073-012-2 |
| SYBR PCR Master mix | Bio-Rad | Cat# 1725124 |
| TRIzol Reagent | Thermo Fisher | Cat# 15596026 |
| Ruxolitinib | Abcam | Cat# AB-ab141356-25-B |
| GKA | Pfizer | Cat# HY-108328 |
| Diazoxide | Sigma-Aldrich | Cat# D9035-250MG |
| Nifedipine | Sigma-Aldrich | Cat# N7634-1G |
| IFNa | BioLegend | Cat# BLG-752804 |
| ExtrAvidin-Peroxidase | Sigma-Aldrich | Cat# E2886 |
| DAB Quanto | Thermo Scientific | Cat# TA-125-HDX |
| Hematoxylin | Sigma-Aldrich |  |
| Zombie Violet™ Dye | Biolegend | Cat# 77477 |
| Perm/Wash™ Perm/Wash Buffer | BD Biosciences | Cat# 54723 |
| Cytofix/Cytoperm™ Fixation and Permeabilization Solution | BD Biosciences | Cat# 554722 |
| Droplet Generation Oil for Probes | Bio-Rad | Cat# D9161172A |
| ddPCR Supermix for Probes | Bio-Rad | Cat# 1863024 |
| Critical commercial assays |  |  |
| Direct-Zol™ RNA-MicroPrep w/ Zymo-Spin™ IC Columns | Zymo | Cat# R2062 |
| qScript cDNA synthesis kit | Quanta Biosciences | Cat# 95047-500 |
| SMARTer Stranded Total RNA-Seq Kit v3 – Pico Input Mammalian | Takara Bio | Cat# 634485 |
| Experimental models: Organisms/strains |  |  |
| Mouse: *Adar*<sup>fl/fl</sup> | (8) | NA |
| Mouse: *MIP-CreER*<sup>TM</sup> | (51) | NA |
| Mouse: *ROSA26-LSL-EYFP* | Jackson Laboratory | B6.129X1-<br>*Gt(ROSA)26Sor*<sup>tm1(EYFP)Cos/J</sup> |
| Software and algorithms |  |  |
| Prism (version 8.0.2) | GraphPad | https://www.graphpad.com/scientific-software/prism/ |
| ImageJ (version 1.8.0) | NIH | http://imagej.nih.gov/ij |
| GSEA |  | https://www.gsea-msigdb.org/gsea/index.jsp |
| FastQC (version 0.11.8) | Babraham Institute | http://www.bioinformatics.babraham.ac.uk/projects/fastqc/ |
| UMI-tools (versions 1.1.1 & 1.1.2) |  | https://umi-tools.readthedocs.io/en/latest/ |
| seqtk trimfq |  | https://github.com/lh3/seqtk |
| STAR (version 2.7.3a) |  | https://github.com/alexdobin/STAR |
| htseq-count script (version 2.0.1) |  | https://htseq.readthedocs.io/en/master/htseqcount.html |
| DeSeq2 (version 1.28.1) |  | https://github.com/mikelove/DESeq2 |
| Other |  |  |
| Collagenase P | Sigma-Aldrich | Cat# 11213873001 |
| RPMI 1640 | Gibco | Cat# 31870025 |
| RPMI 1640 no glucose | Sartorius | Cat# 01-101-1A |
| Histopaque-1119 (Ficoll 1.119g/mL) | Sigma-Aldrich | Cat# 111910-100ML |
| Histopaque-1077(Ficoll 1.079 g/mL) | Sigma-Aldrich | Cat# 10771-100ML |

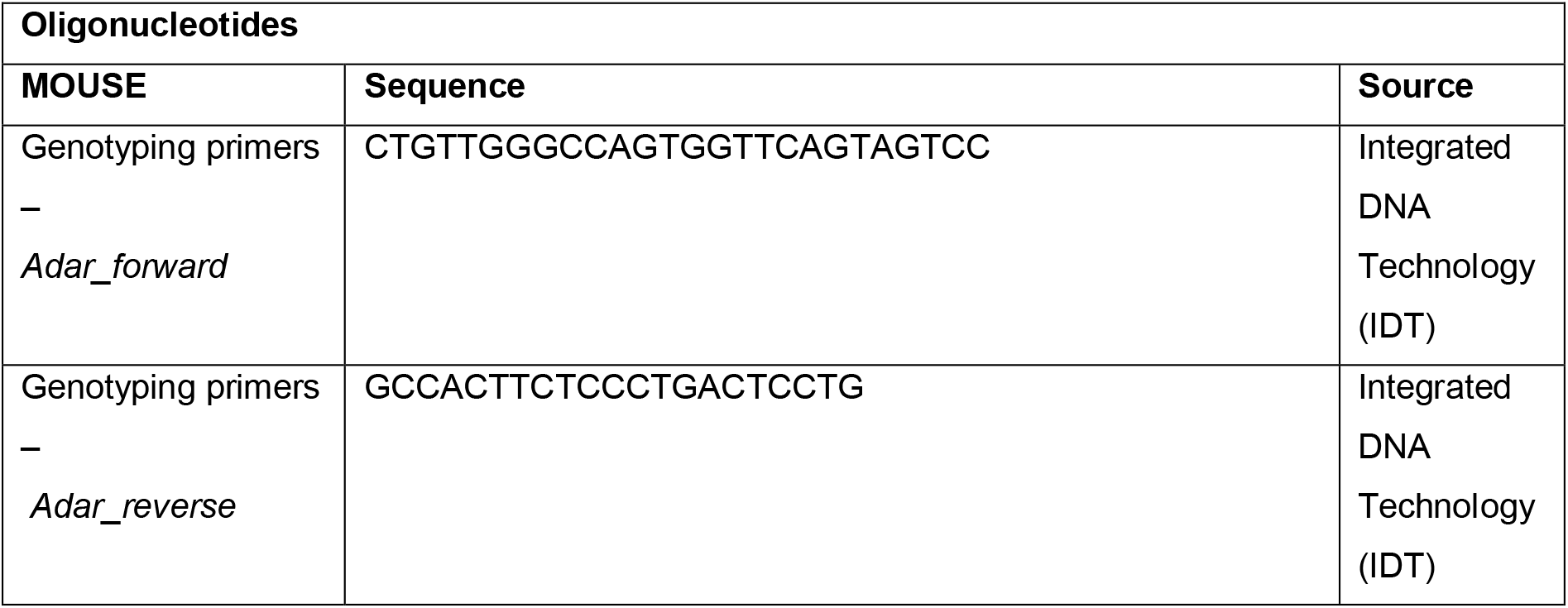
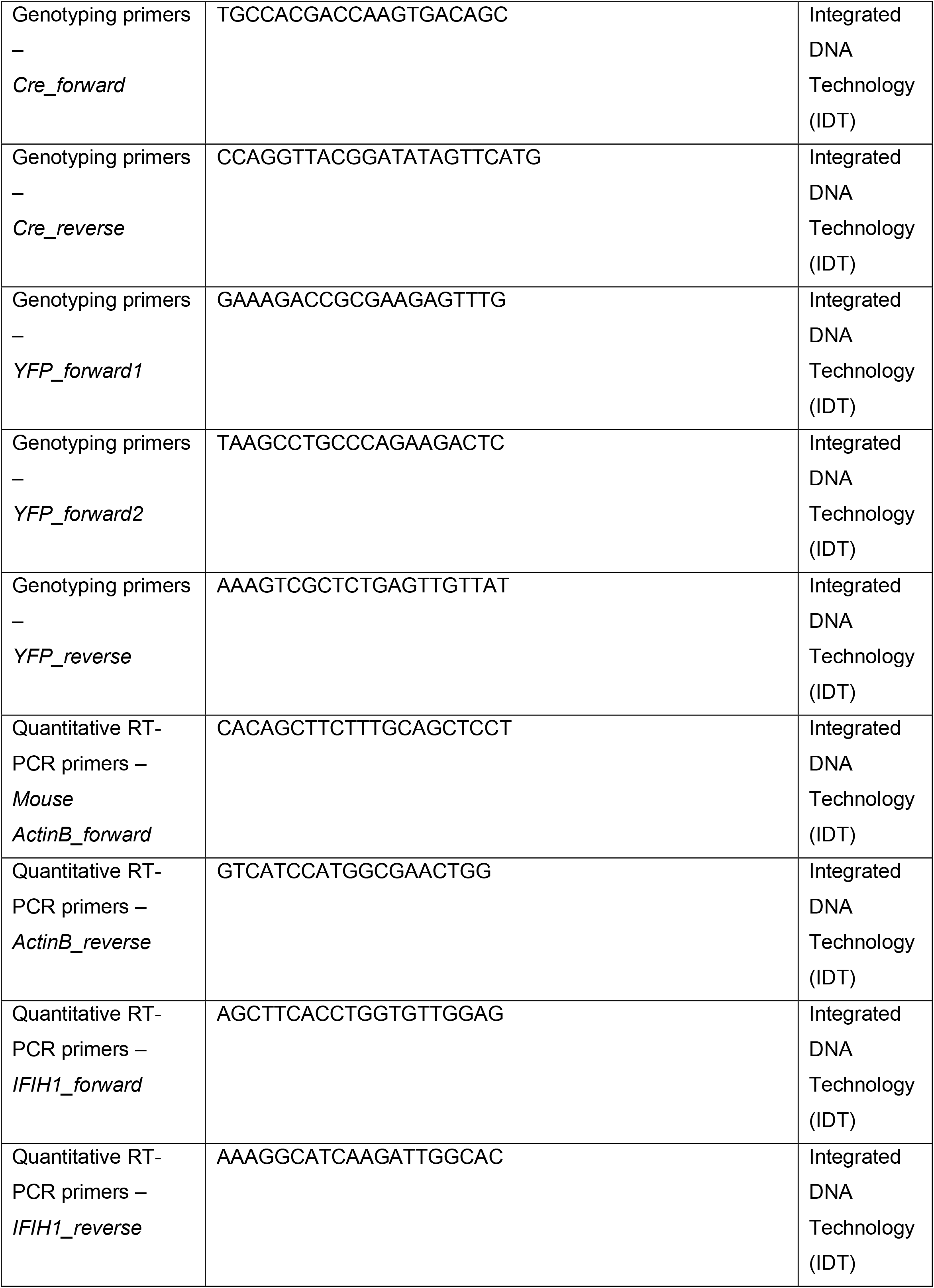
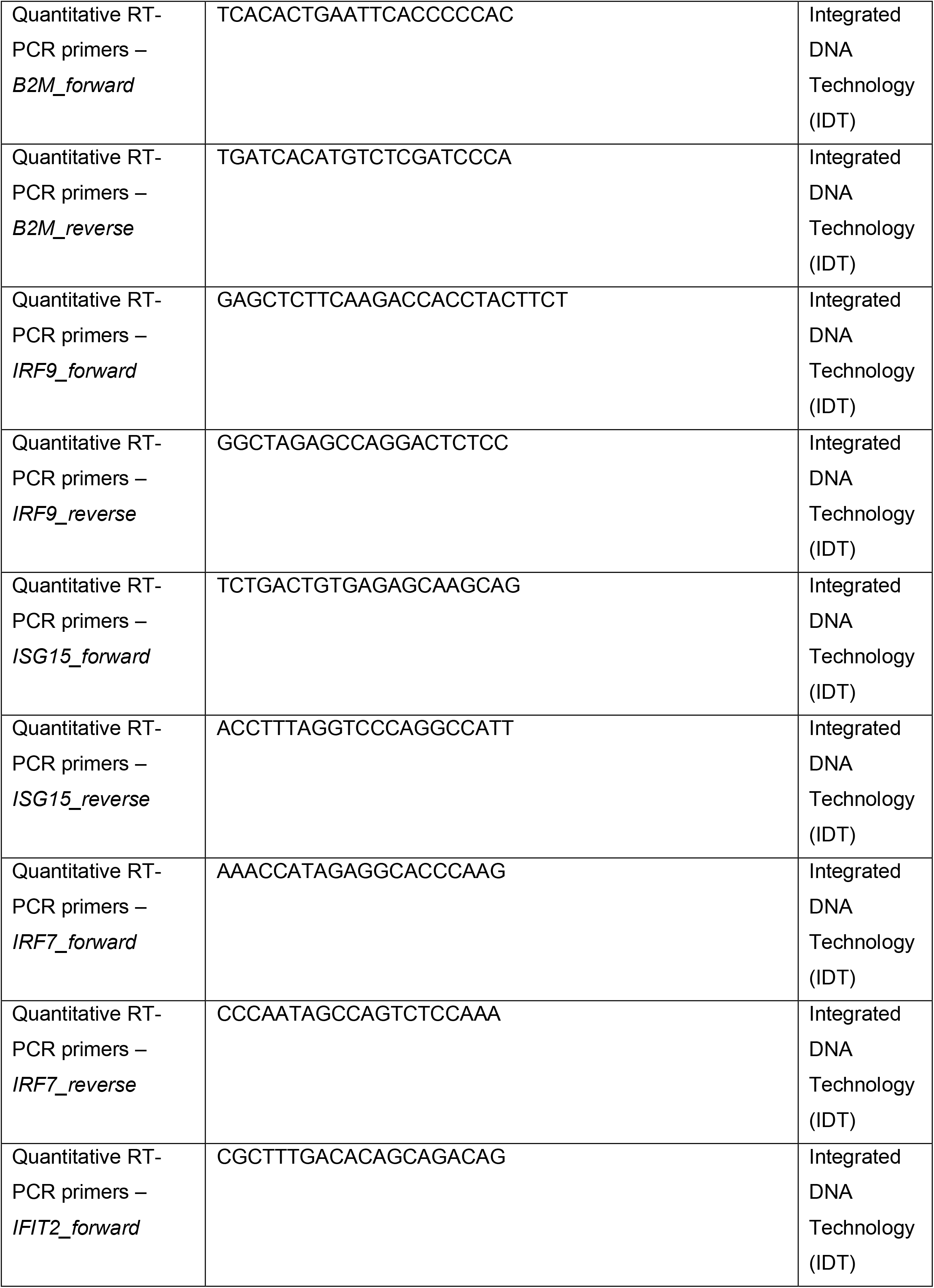
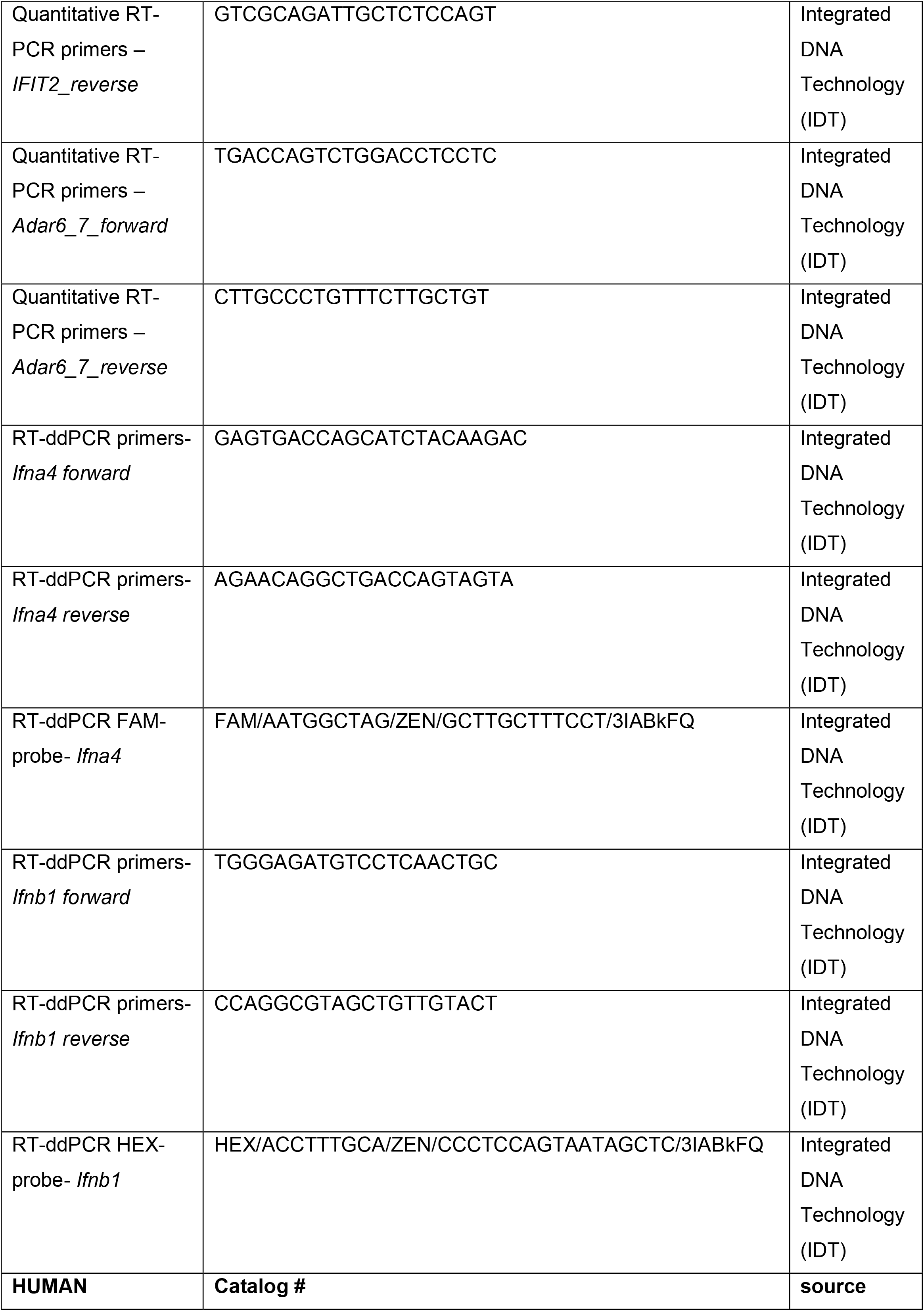
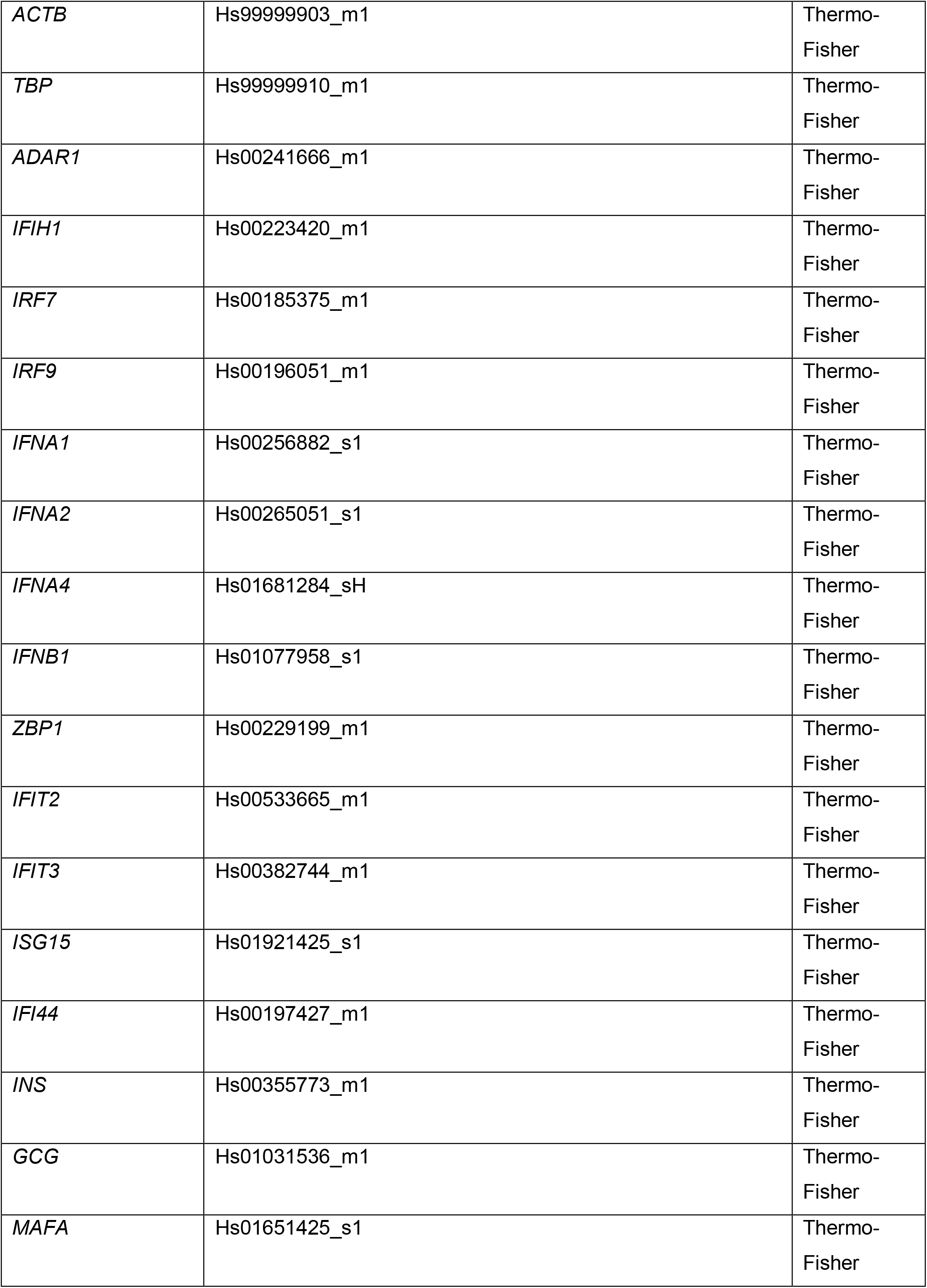
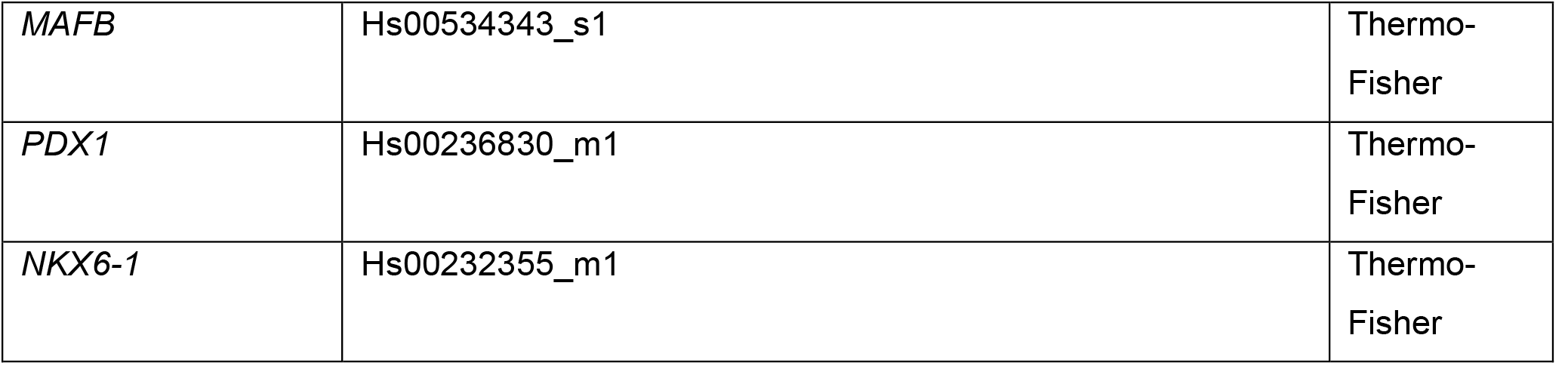

**Fig. S1.**
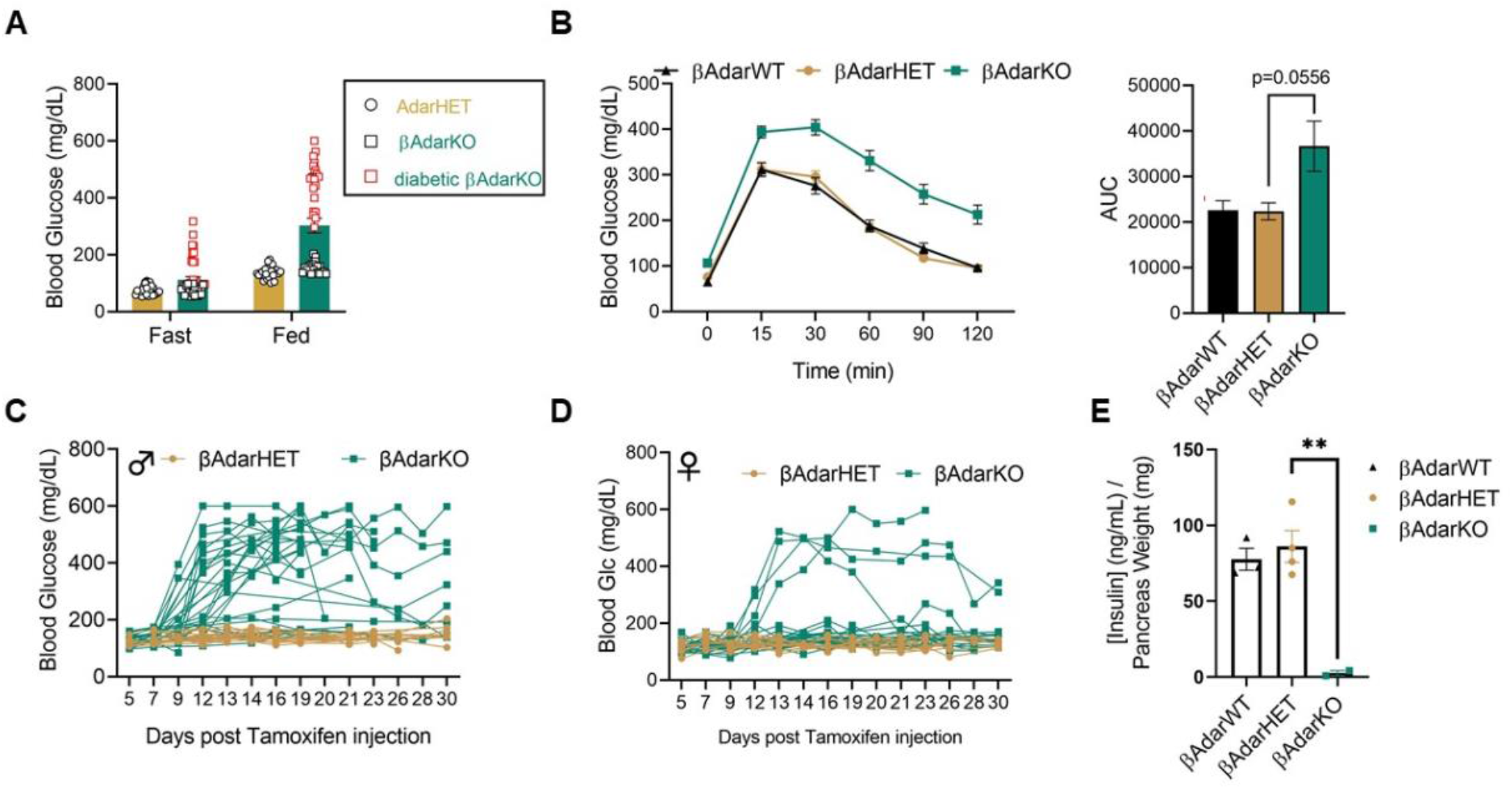
Glycemic phenotype in βAdar-mutant mice. (**A**) Fasting and random (fed) blood glucose levels were measured in 1-month-old βAdarHET (n=25) and β-AdarKO (n=48) mice 14-15 days after tamoxifen injection. Red squares represent diabetic β-AdarKO mice. (**B**) β-AdarKO mice show glucose intolerance. 1-month-old βAdarWT (n=17), βAdarHET (n=25) and β-AdarKO (n=48) mice were injected with tamoxifen and intraperitoneal glucose tolerance test (IP-GTT) was performed 15 days later. Right bar plot represents area under the glucose curves (AUC). Data are presented as mean ± SEM. (**C, D**) βAdarHET (19 males and 17 females) and βAdarKO (27 males and 25 females) were monitored for fed glucose from day 5 after Tamoxifen injection. Results show a more pronounced diabetic phenotype in βAdarKO male (**C**) compared to βAdarKO female (**D**) mice. (**E**) Pancreatic insulin content is reduced in diabetic βAdarKO mice. 1-month-old βAdarWT, βAdarHET and β-AdarKO male mice were injected with tamoxifen and their pancreas harvested 16 days later. Pancreas insulin content was measured by ELISA and normalized to pancreas weight.

**Fig. S2:**
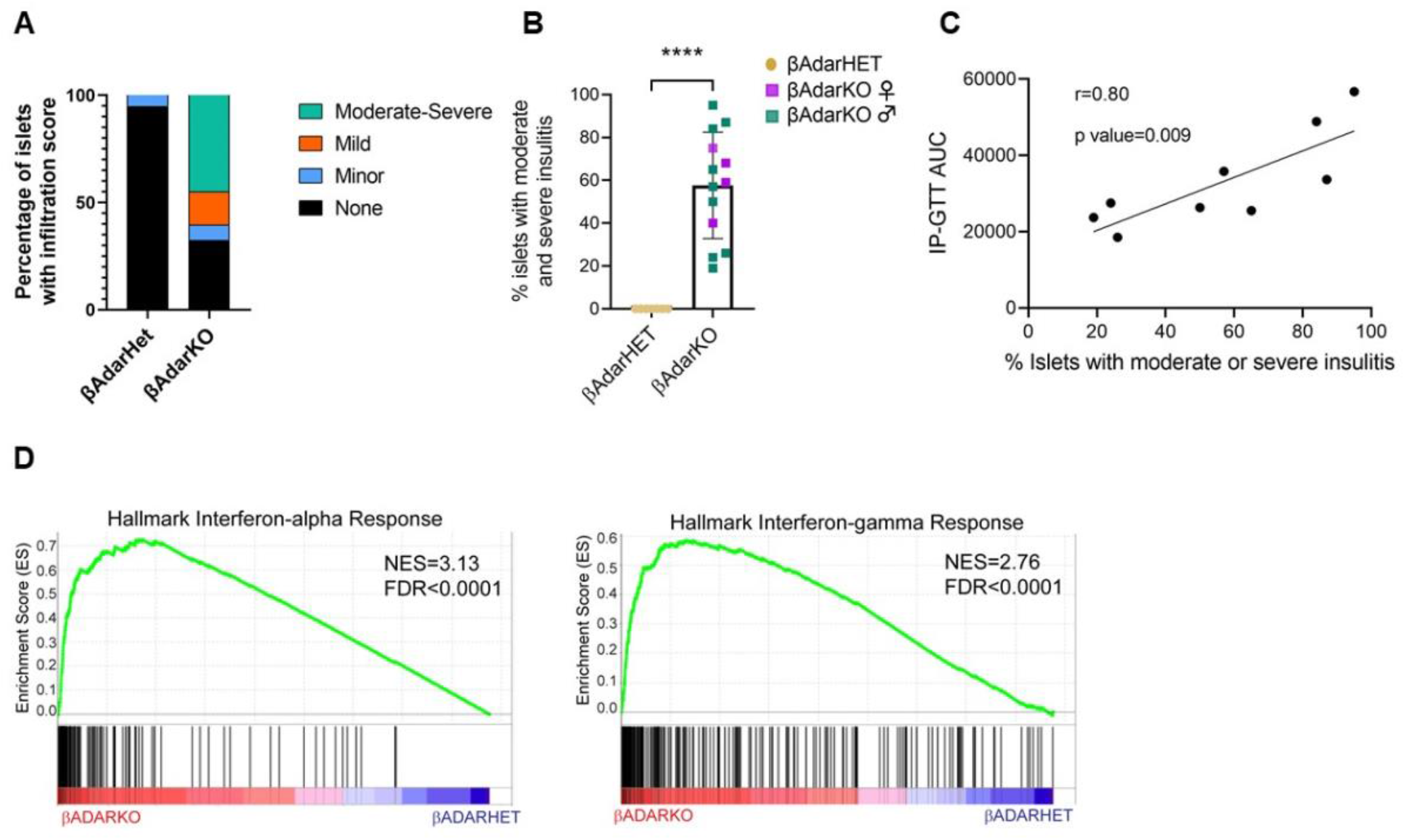
Insulitis in βAdarKO mice. (**A**) Pooled insulitis scores (see Figure 2A and methods) of pancreatic sections from β-AdarHET (n=4) and β-AdarKO (n=13) 1-month-old mice 12-14 days after tamoxifen injection. (**B**) Insulitis is not affected by mice gender. Insulitis scores of β-AdarKO shown in Figure 2A reveal similar extent of immune cell infiltration in males and females. (**C**) Correlation of impaired glucose tolerance to extent of insulitis. IP-GTT was performed on βAdarKO male mice (n=9) 9 days after Tamoxifen injection and 3 days later pancreatic islet insulitis score was determined as described in Figure 2A. The values of the area under the GTT curve (IP-GTT AUC) were plotted versus insulitis score. (**D**) GSEA plots showing that genes of the interferon-alpha and gamma response are highly enriched among the genes upregulated in AdarKO versus AdarHET.

**Fig. S3:**
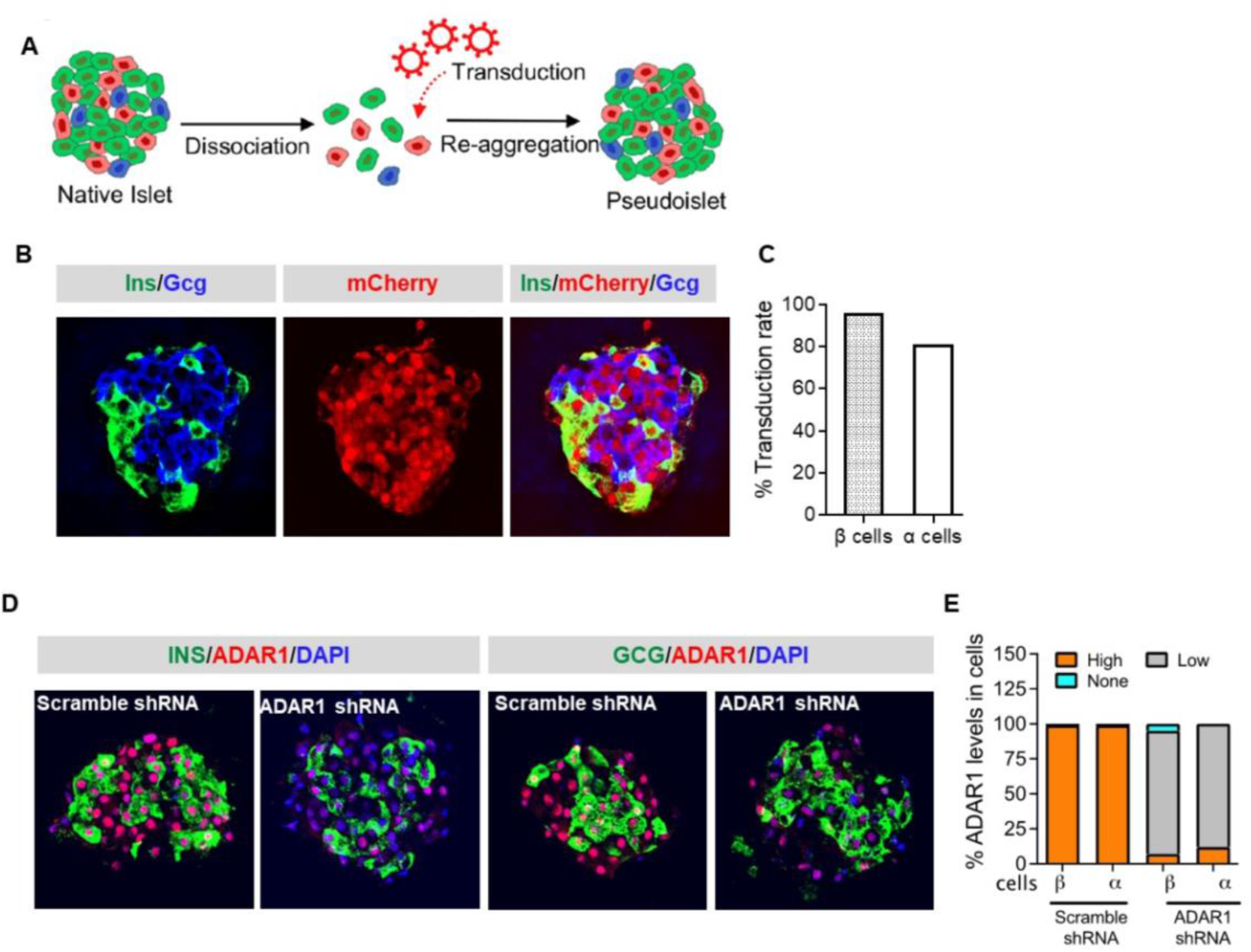
Adar1 knockdown in human islets. **(A)** Schematic of experiment: human islet dissociation, infection and reaggregation to form pseudoislets. **(B)** To assess the efficiency of transduction, pseudoislets were immobilized in 3-D collagen gels and sectioned for immunofluorescence staining as described (*52*). The cells were labeled with insulin (green), mChery (red), and glucagon (blue) antibodies. Images were acquired with a confocal laser-scanning microscope. The images are representative of pseudoislets from both groups group. **(C)** Quantification of virus transduction rate. In scrambled shRNA pseudoislets, 96% of beta cells and 81% alpha cells were mCherry positive. A similar % of mCherry positive cells were seen in shADAR and scrambled shRNA pseudoislets. **(D)** Representative images of pseudoislets labeled with insulin or glucagon (green), ADAR1 (red), and DAPI (blue) in scrambled shRNA or ADAR1 shRNA transduced pseudoislets. **(E)** % OF ADAR1 expression levels in beta and alpha cells in pseudoislets. Brown = high level, grey=low level, cyan=no ADAR1.

**Fig. S4.**
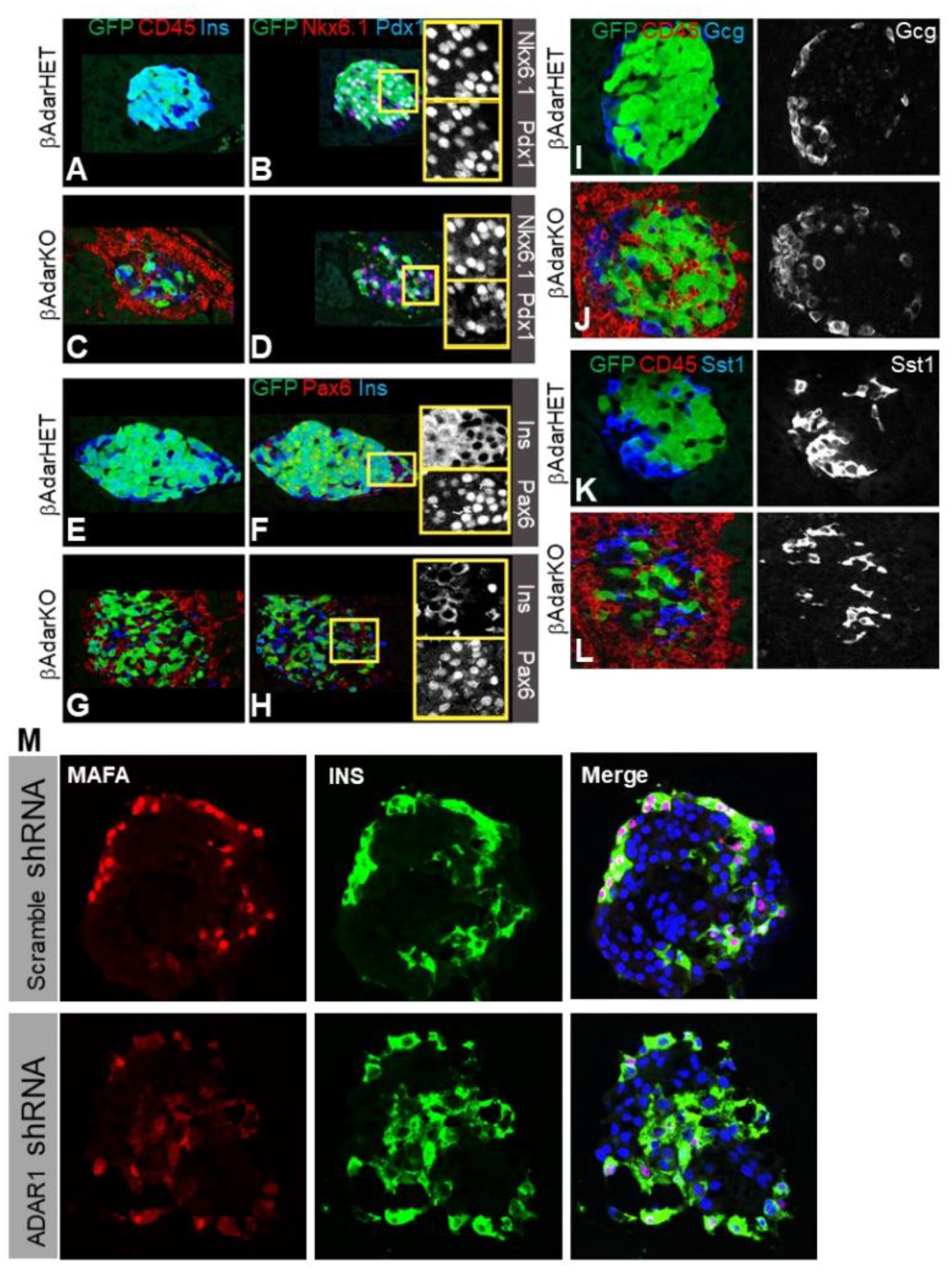
Expression of beta-cell transcription factors in βAdarKO mice and in ADAR1-deficient human islets. **(A-H)** Persistence of PDX1, NKX6.1 and PAX6 levels in inflamed islets of βAdarKO mice. Pancreatic sections from non-diabetic βAdarHET(**A, B, E, F**) and βAdarKO mice (**C, D, G, H**), 12 days after Tamoxifen injection were immunostained for GFP, CD45 and Insulin (**A, C, E, G**) and their serial sections were stained for GFP, PDX1 and NKX6.1 (**B, D**) and GFP, PAX6 and Insulin (**F, H**). Representative micrographs show that insulitis does not affect the transcription factor levels in beta-cells. (**I-L**) Alpha and delta cells are spared in the face of insulitis in βAdarKO mice. Pancreatic sections from non-diabetic βAdarKO and control βAdarHET mice, 12 days after Tamoxifen injection were immunostained for GFP, CD45 and Glucagon (**I-J**) and GFP, CD45 and Somatostatin (**K-L**). (**M**) Reduced MAFA levels in cultured human pseudoislets with ADAR1 knockdown. The cells were labeled with MAFA (red), insulin (green), and DAPI (blue).

**Fig. S5:**
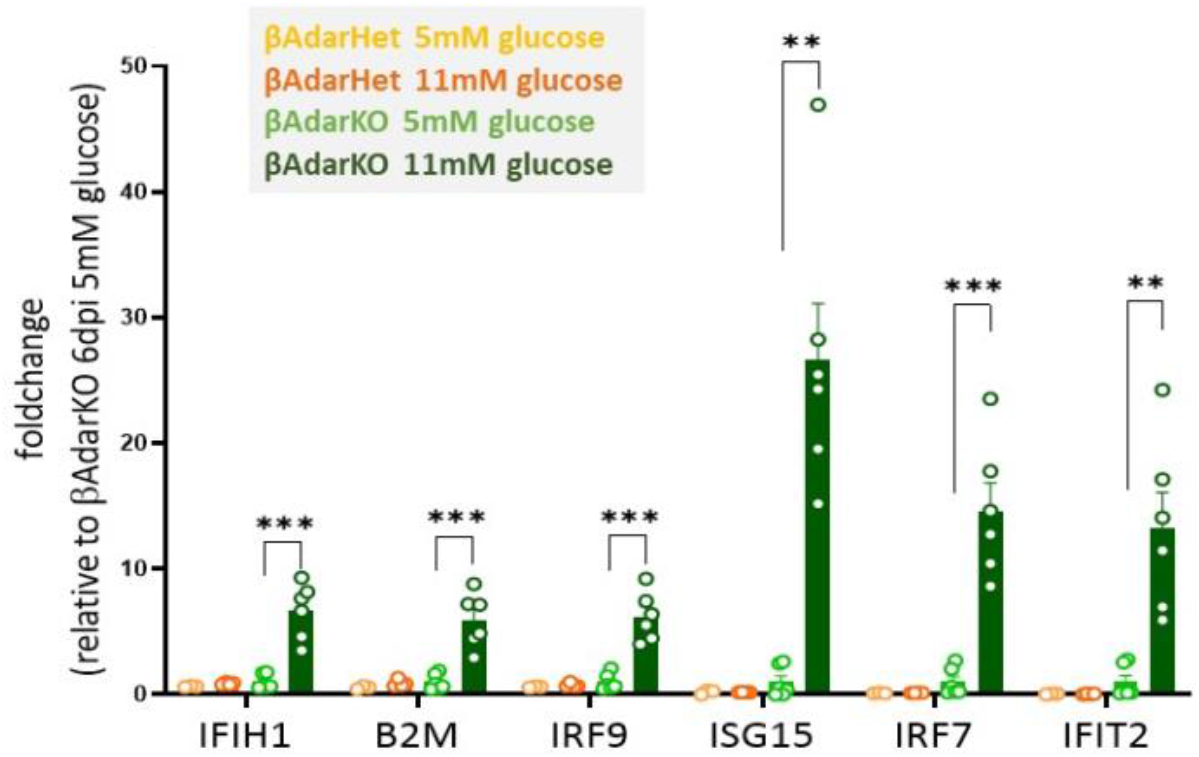
Low glucose reduces the interferon response in in islets from βAdarKO mice. Islets from βAdarHET and βAdarKO mice were isolated 3 days after tamoxifen injection and cultured for 3 days in RPMI medium supplemented with 5mM or 11mM glucose. ISG expression in whole islets was assayed by RT-qPCR and normalized to Actb expression. n>=6. *p<0.05, **p<0.01; Student’s unpaired two-tailed t-test; error bars indicate mean ± SEM.

**Fig. S6:**
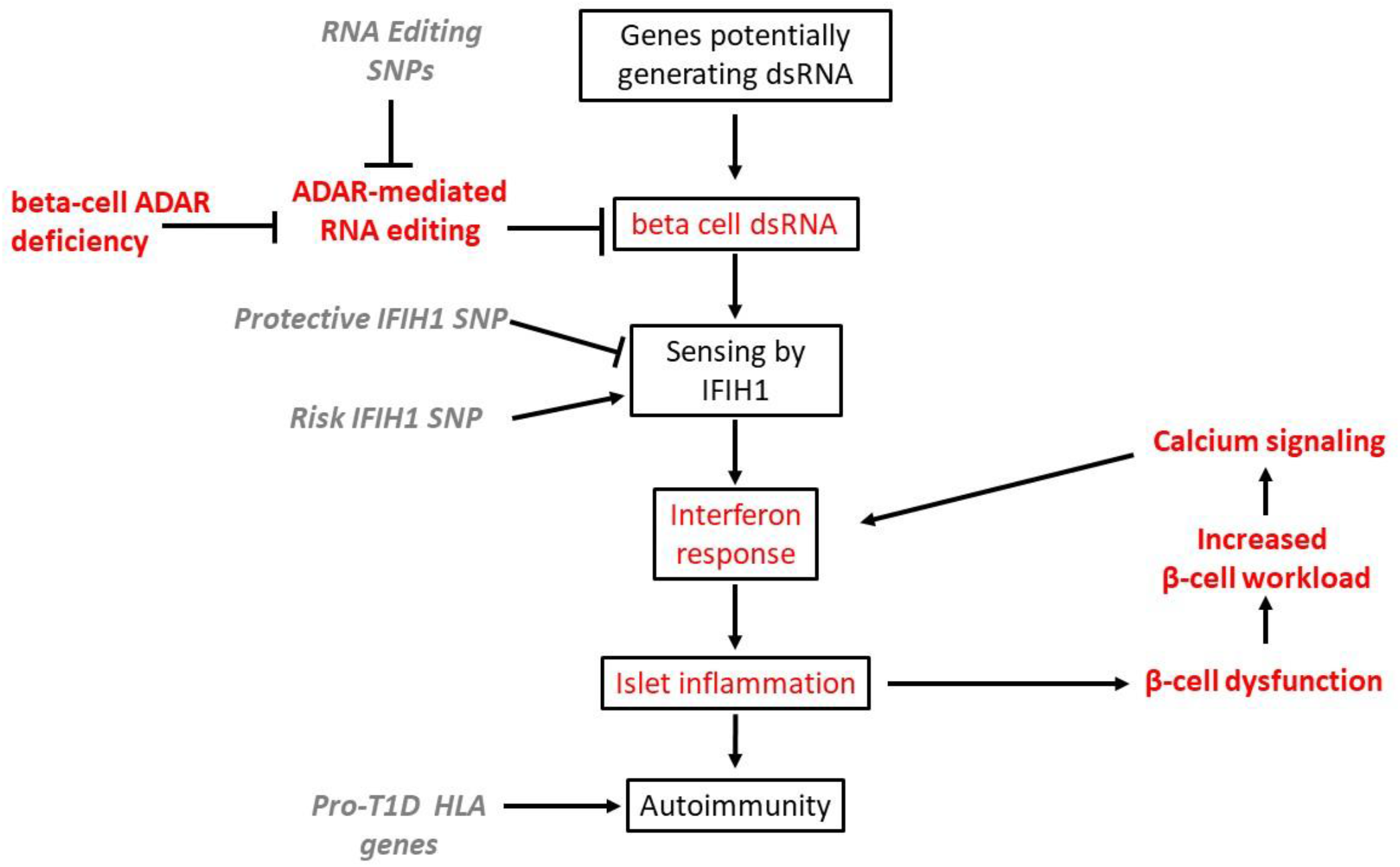
Schematic model for the involvement of defective RNA editing in the pathogenesis of T1D. This model is supported by our results (in red) and published human genetic evidence (in grey italics)

**Fig. S7:**
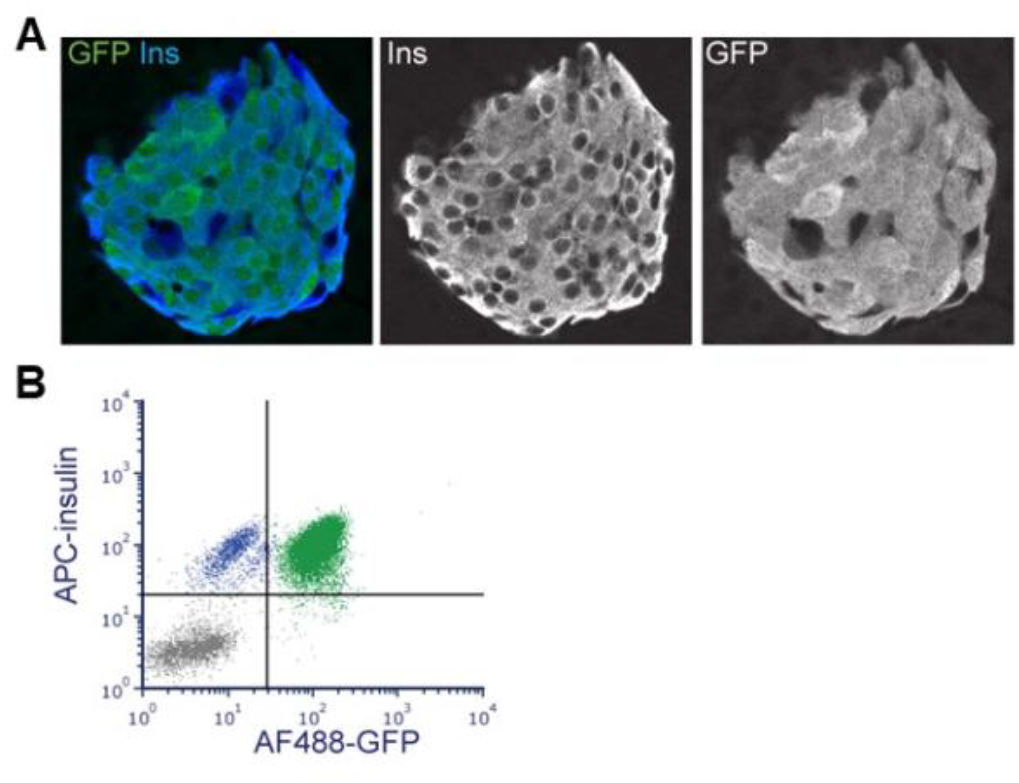
GFP expression induced by MipCre-ER is a faithful reporter of beta-cells. Pancreatic sections (**A**) and isolated islets (**B**) from 1-month-old βAdarHET mice 12 days after injection of tamoxifen were immunostained for GFP and insulin. (**A**) Representative islet from a pancreatic section and (**B**) FACS analysis of dissociated immunostained islets show perfect overlap of insulin and GFP staining.

**Fig. S8:**
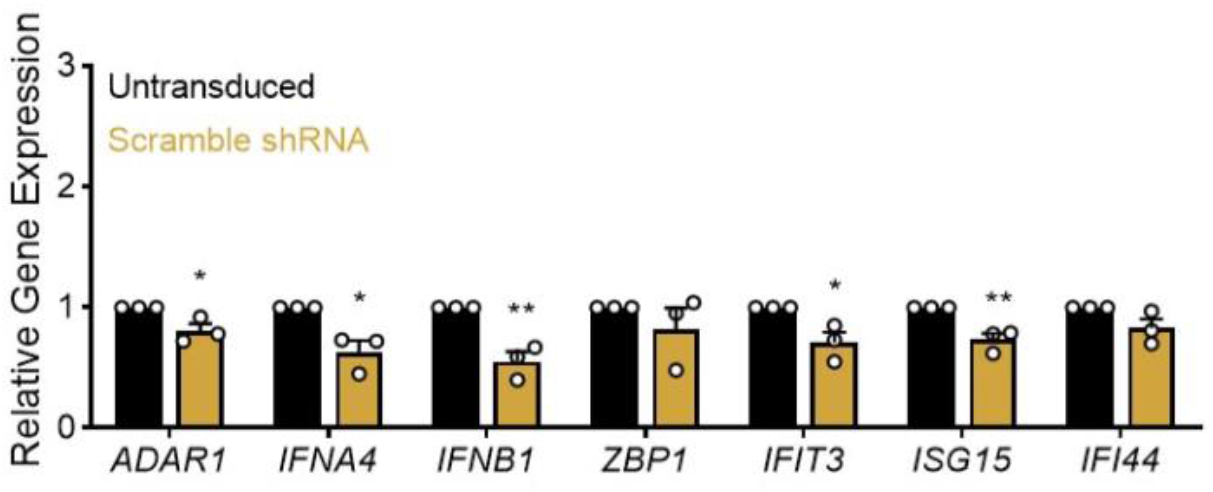
Adenoviral infection does not induce type I IFNs in human islets. RNA isolated from untransduced pseudoislets and scramble virus transduced pseudoislets was assessed by RT-PCR. The mRNA for ADAR1, IFN, and IFN-induced genes was slightly lower in scramble-transduced islets. * p<0.05; ** p<0.01

**Table S1:**
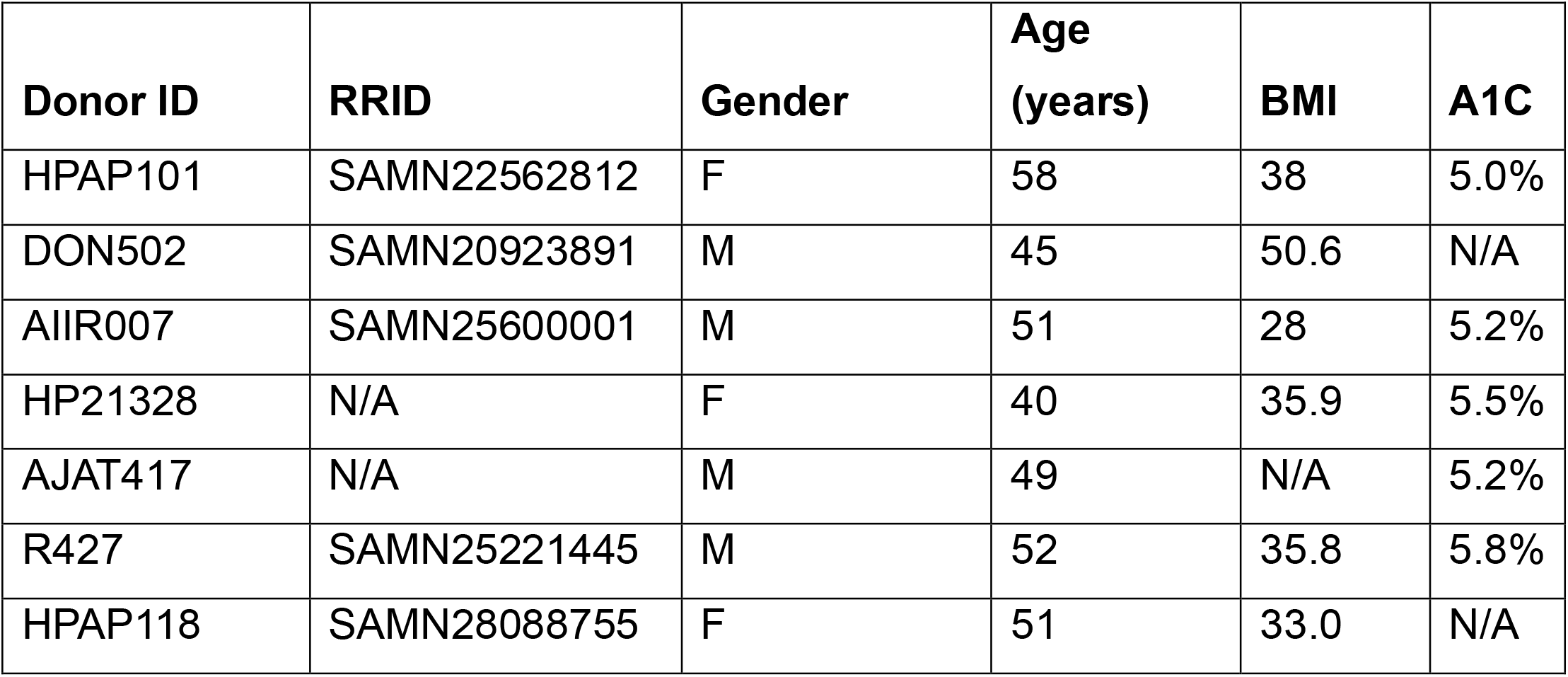
Donor Information.

